# Widespread detoxifying NO reductases impart a distinct isotopic fingerprint on N_2_O under anoxia

**DOI:** 10.1101/2023.10.13.562248

**Authors:** Renée Z. Wang, Zachery R. Lonergan, Steven A. Wilbert, John M. Eiler, Dianne K. Newman

## Abstract

Nitrous oxide (N_2_O), a potent greenhouse gas, can be generated by compositionally complex microbial populations in diverse contexts. Accurately tracking the dominant biological sources of N_2_O has the potential to improve our understanding of N_2_O fluxes from soils as well as inform the diagnosis of human infections. Isotopic “Site Preference” (SP) values have been used towards this end, as bacterial and fungal nitric oxide reductases produce N_2_O with different isotopic fingerprints. Here we show that flavohemoglobin, a hitherto biogeochemically neglected yet widely distributed detoxifying bacterial NO reductase, imparts a distinct SP value onto N_2_O under anoxic conditions that correlates with typical environmental N_2_O SP measurements. We suggest a new framework to guide the attribution of N_2_O biological sources in nature and disease.

**One-Sentence Summary:** Detoxifying nitric oxide reductases impart a distinct isotopic biosignature on nitrous oxide.

## Main Text

Nitrous oxide (N_2_O) is a ubiquitous metabolite present in myriad environments ranging from soils, marine and freshwater systems, and the atmosphere to the human body. Because N_2_O can be produced and consumed by multiple microbial nitrogen-cycling processes (*1*), tracking its sources and fates is challenging. One motivation to do so springs from the fact that N_2_O is a potent greenhouse gas, whose current atmospheric concentration is more than 20% compared to preindustrial levels (*2*); a better understanding of N_2_O sources could help facilitate mitigation efforts. Analogously, because N_2_O has been measured in chronic pulmonary infections (*3*), clarity on which pathogens are metabolically active in disease contexts could inform treatment strategies (*4*).

An intramolecular isotopic fingerprint called “Site Preference” (SP), which measures the relative enrichment of natural abundance ^15^N in the central (□) versus terminal (□) nitrogen position in N_2_O (Fig. 1A; (*5*)) may be applied for such purposes. Unlike traditional natural abundance isotopic measurements of the total ^15^N in the bulk molecule (*6*), SP does not rely on the isotopic composition of the source substrate but instead reflects the reaction mechanism (*7*), making it a potentially powerful tool to disentangle N_2_O sources in different contexts.

**Fig. 1.**
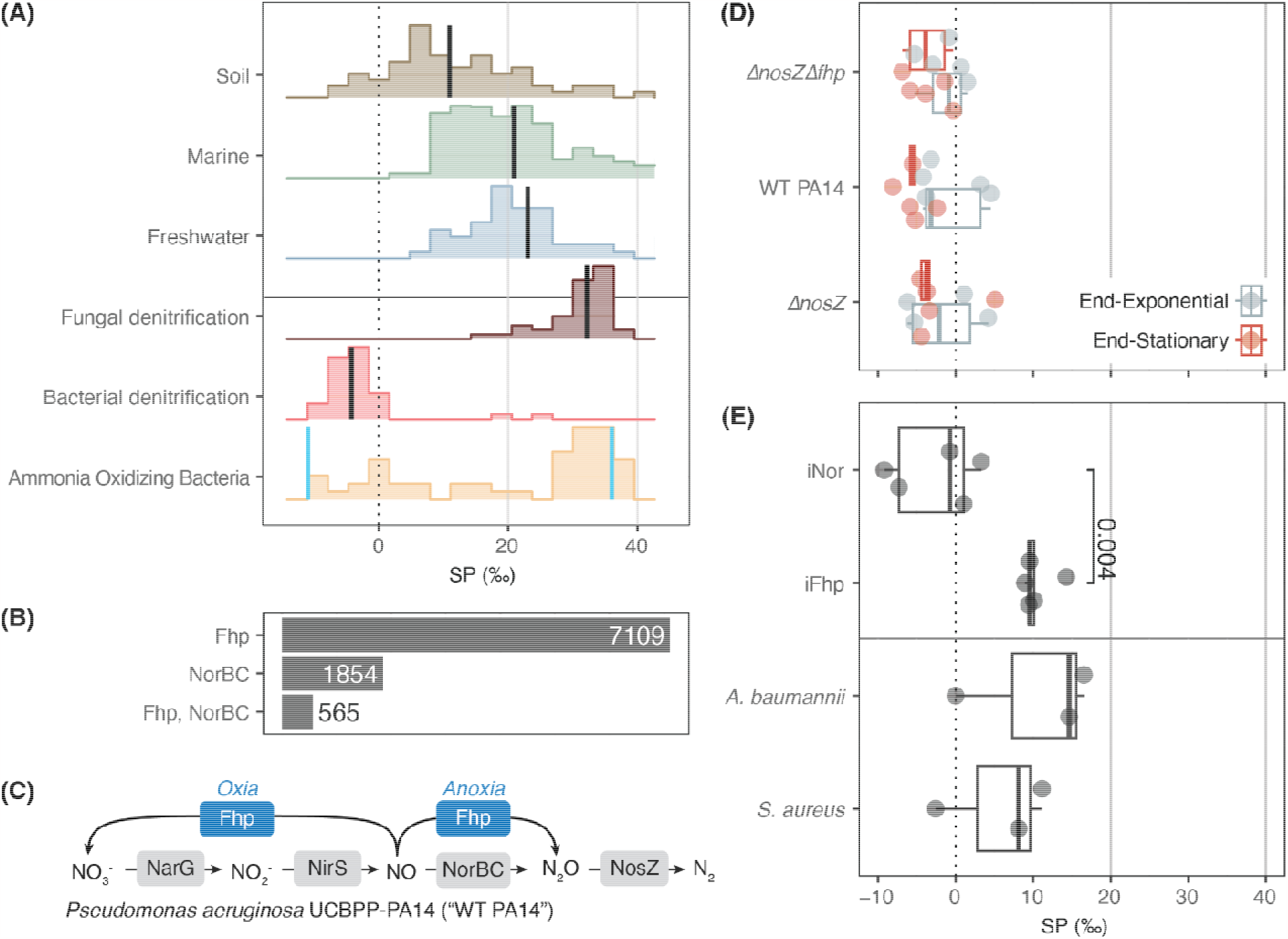
N_2_O production via NO detoxification under anoxic conditions may explain environmental SP values. **(A)** Measured *in situ* SP values for environmental (Soil, Marine, Freshwater) vs. *in vitro* measurements of biogenic end-members (Bacterial and Fungal Denitrification, Ammonia Oxidizing Bacteria (AOB)); black line shows median; blue lines show end-member values for AOB (*8*). Histogram height is normalized to each category; see Fig. S11 for outlier values and more detail. **(B)** Number of bacterial genomes hits at the phylum level for flavohemoglobin protein (Fhp) and nitrous oxide reductase (NorBC) alone or in combination from Annotree (*11*); minimum amino acid sequence similarity of 30% was used. See Fig. S2, Tables S1-S4 for phylogenetic distribution. **(C)** Relevant N-oxide pathways of *Pseudomonas aeruginosa* UCBPP-PA14 (*Pa*), the model organism used in this study. *Pa* possesses the full denitrification pathway as well as Fhp. **(D)** SP of N_2_O produced by *Pa* and mutant strains with *fhp* and/or *nosZ* genes deleted (Δ*nosZ*Δ*fhp*; Δ*nosZ*) in denitrifying conditions; see Fig. S2 for more detail. **(E)** of *Pa* strains with rhamnose-induced expression of *norBC* (iNor) or *fhp* (iFhp) alone as well as *Acinetobacter baumannii* and *Staphylococcus aureus*, which only have Fhp. *P* value was calculated via Welch’s t-test. Each data point represents an individual biological replicate in (D) and (E).

The median values of *in situ* SP measurements where microbes are present are 10.9, 20.9 and 23.0 per mille (‰) for soils, marine and freshwater systems, respectively (Fig. 1A). These values are bounded by the median values of *in vitro*, pure culture studies of N_2_O-producing biogenic end-members like bacterial and fungal denitrifiers as well as ammonia-oxidizing bacteria (AOB; Fig. 1A). Bacterial and fungal denitrifiers are thought to represent two extremes of SP values for N_2_O producers with median SP values of -4.3 and 32.2‰ respectively (Fig. 1A), which are assumed to reflect the activity of dissimilatory Nitric Oxide Reductases (NOR); in AOBs, the SP varies between roughly -11 and 36‰ due to multiple dissimilatory N_2_O formation pathways (*8*). Because the vast majority of *in situ* environmental observations lie between end-member values for bacterial and fungal NORs and AOBs, the SP values of biogenic N_2_O produced in the environment have been rationalized by mixing of these end-members, which assume the activity of catabolic pathways that are tied to microbial growth.

However, an entire other class of enzymes exists that produce N_2_O as a consequence of nitric oxide (NO) detoxification and not for energy-conservation (*9*). Flavohemoglobin proteins (e.g. Fhp/Hmp/Yhb–henceforth referred to as “Fhp”) are phylogenetically widespread and protect against nitrosative stress in bacteria and yeast (*10*). Members of this family are roughly four times more abundant than NORs in annotated bacterial genomes (Fig. 1B, Fig. S1, Tables S1-3; 7109 vs. 1854 genome hits at the phylum level for Fhp vs. NorBC using 30% minimum amino acid sequence similarity (*11*)). While their ability to oxidize NO to nitrate (NO_3_^-^) under oxic conditions is well known, their ability to reduce NO to N_2_O under anoxic conditions has received less attention (*10, 12*). Given that bacterial denitrifiers commonly possess both Fhp and NOR (Fig. 1B and Table S1), we hypothesized that Fhp might play a role in N_2_O emissions and set out to determine whether it imparts a SP onto N_2_O distinct from that of bacterial or fungal NORs.

## Overall SP values reflect NOR during denitrification

To compare the SP of Fhp to NOR in a whole cell context (*in vivo*), we used the model bacterial denitrifier, *Pseudomonas aeruginosa* UCBPP-PA14 (*Pa*, Fig. 1C). Because this organism is genetically tractable, it provides a means to study the cellular processes of interest in a controlled manner (Table 1). To determine SP values under denitrifying conditions, wild type (WT) *Pa*, Δ*nosZ* and Δ*nosZ*Δ*fhp* – strains with deletions of the nitrous oxide reductase (NOS) gene, *nosZ* (*PA14_20200*) and/or *fhp* (*PA14_29640*) – were grown anaerobically in defined medium batch cultures and sampled at late exponential and late stationary growth phase (Table 2, Fig. S2 and Methods). N_2_O was cryogenically distilled and analyzed for nitrogen and oxygen isotopes on the Thermo Scientific Ultra High-Resolution Isotope Ratio Mass Spectrometer (HR-IRMS; (*13*); Methods). All isotope data is reported in the delta (δ) notation in units of per mille (‰) where δ^15^N = [(^15^N/^14^N)_sample_ / (^15^N/^14^N)_reference_ - 1]*1000 and SP = δ^15^N^□^ - δ^15^N^□^. Values are reported relative to the international reference of AIR for nitrogen; see Methods for more detail.

**Table 1.** Strains studied. The SP of N_2_O produced by five strains of *Pseudomonas aeruginosa* (WT *Pa*, Δ*nosZ*, Δ*nosZ*Δ*fhp, iFhp, iNOR*) and two wild-type strains of *Staphylococcus aureus* and *Acinetobacter baumannii* were measured. See Materials and Methods for further detail. *S. aureus* and *A. baumannii* were both kindly provided by Eric Skaar, Vanderbilt University Medical Center.

| Name | Strain Description | Fhp? | Nor? | Source |
| --- | --- | --- | --- | --- |
| WT *Pa* | Wild-type *Pseudomonas aeruginosa* UCBPP-PA14 | Yes | Yes | Lab Collection |
| Δ*nosZ* | Deletion of nitrous oxide reductase gene (*nosZ*, PA14_20200) from WT *Pa* | Yes | Yes | This study |
| Δ*nosZ*Δ*fhp* | Deletion of *nosZ* and flavohemoglobin protein (*fhp*, PA14_29640) from WT *Pa* | Yes | No | This study |
| iFhp | Rhamnose-induced expression of *fhp* integrated into the chromosome of WT *Pa* with deletion of native *norBC*, *fhp*, and *nosZ*. | Yes | No | This study |
| iNOR | Rhamnose-induced expression of the nitric oxide reductase operon, *norBCD* (PA14_16810, PA14_16830, PA14_06840), integrated into the *att* neutral chromosomal site of *Pa* with deletion of native nitrate reductase (*narGHJI*; PA14_13780-13830), nitrite reductase (*nirS*; PA14_06750), *norBC*, *nosZ*, and *fhp*. | No | Yes | This study |
| *S. aureus* | Wild-type *Staphylococcus aureus* USA300 LAC | Yes | No | Gift |
| *A. baumannii* | Wild-type *Acinetobacter baumannii* ATCC 17978 | Yes | No | Gift |

**Table 2.**
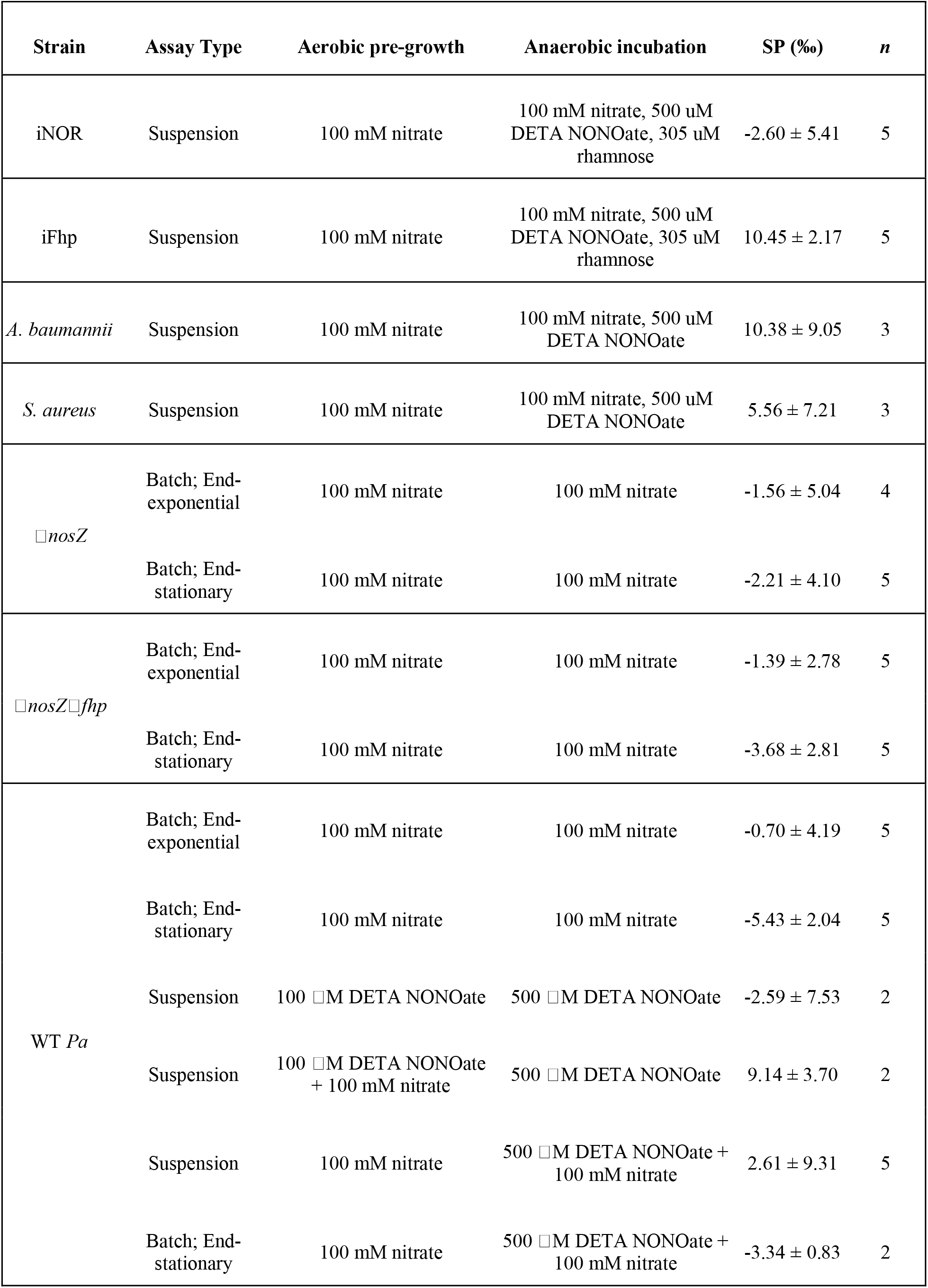
Culturing conditions and SP results. All strains were grown in aerobic pre-growths before being resuspended in fresh media and anoxically incubated for headspace sampling as batch culture or suspension assays (Fig. S12); nitrate and/or DETA NONOate (C_4_H_13_N_5_O_2_) was supplemented to provide endogenous vs. exogenous NO respectively. See Methods for more detail. SP values (mean ± s.d.) of *n* biological replicates; see Supplemental for full data table.

| Strain | Assay Type | Aerobic pre-growth | Anaerobic incubation | SP (‰) | *n* |
| --- | --- | --- | --- | --- | --- |
| iNOR | Suspension | 100 mM nitrate | 100 mM nitrate, 500 uM DETA NONOate, 305 uM rhamnose | $-2.60 \pm 5.41$ | 5 |
| iFhp | Suspension | 100 mM nitrate | 100 mM nitrate, 500 uM DETA NONOate, 305 uM rhamnose | $10.45 \pm 2.17$ | 5 |
| *A. baumannii* | Suspension | 100 mM nitrate | 100 mM nitrate, 500 uM DETA NONOate | $10.38 \pm 9.05$ | 3 |
| *S. aureus* | Suspension | 100 mM nitrate | 100 mM nitrate, 500 uM DETA NONOate | $5.56 \pm 7.21$ | 3 |
| □*nosZ* | Batch; End-exponential | 100 mM nitrate | 100 mM nitrate | $-1.56 \pm 5.04$ | 4 |
| | Batch; End-stationary | 100 mM nitrate | 100 mM nitrate | $-2.21 \pm 4.10$ | 5 |
| □*nosZ*□*fhp* | Batch; End-exponential | 100 mM nitrate | 100 mM nitrate | $-1.39 \pm 2.78$ | 5 |
| | Batch; End-stationary | 100 mM nitrate | 100 mM nitrate | $-3.68 \pm 2.81$ | 5 |
| WT *Pa* | Batch; End-exponential | 100 mM nitrate | 100 mM nitrate | $-0.70 \pm 4.19$ | 5 |
| | Batch; End-stationary | 100 mM nitrate | 100 mM nitrate | $-5.43 \pm 2.04$ | 5 |
| | Suspension | 100 □M DETA NONOate | 500 □M DETA NONOate | $-2.59 \pm 7.53$ | 2 |
| | Suspension | 100 □M DETA NONOate + 100 mM nitrate | 500 □M DETA NONOate | $9.14 \pm 3.70$ | 2 |
| | Suspension | 100 mM nitrate | 500 □M DETA NONOate + 100 mM nitrate | $2.61 \pm 9.31$ | 5 |
| | Batch; End-stationary | 100 mM nitrate | 500 □M DETA NONOate + 100 mM nitrate | $-3.34 \pm 0.83$ | 2 |

The SP of Δ*nosZ*Δ*fhp* should only reflect NOR, since all other known pathways for N_2_O production and consumption were deleted. The *in vivo* SP of this strain did not vary significantly by growth phase (Welch’s *t*-test, *P*=0.2), and its average value across all growth phases (-2.53 ± 2.90, mean ± s.d. throughout, *n* = 10) was consistent with prior *in vitro* measurements of NOR purified from *Paracoccus denitrificans* ATCC 35512 (-5.9 ± 2.1‰, (*14*)). The SP of the Δ*nosZ*Δ*norBC* strain, which only has *fhp*, was not measured because it did not grow appreciably under these conditions (Fig. S2) presumably due to growth suppression when NO build-up is too high (*15, 16*).

WT *Pa*, which can produce N_2_O through both Fhp and NOR (Fig. 1C), displayed SP values that did not vary significantly from those observed for the Δ*nosZ*Δ*fhp* strain across all growth phases when denitrifying (*P*=0.7). In addition, the SP of WT *Pa* did not vary significantly by growth phase (*P*=0.07). The SP of Δ*nosZ* was also measured because prior studies showed that NOS can increase the SP of the residual N_2_O pool through preferential cleavage of the ^14^N-O vs. ^15^N-O bond in N_2_O (*17, 18*); however, SP values of Δ*nosZ* were similar to Δ*nosZ*Δ*fhp* (*P*=0.7) and did not vary by growth phase (*P*=0.8; Fig. 1D). Therefore, even though Fhp was likely present in all previously measured bacterial denitrifier strains for *in vitro* measurements (Table S2), it does not affect the overall SP value when strains are grown under denitrifying conditions, suggesting that NOR dominates the isotopic signature under these conditions. However, the potential for Fhp to impact the SP of N_2_O under other conditions remained open.

## Fhp has a intermediate, positive SP value compared to bacterial and fungal NORs

To distinguish the SP of Fhp and NOR, we engineered two *Pa* strains possessing only Fhp or NOR that could be induced in the presence of rhamnose; inducible Fhp (“iFhp”) and NOR (“iNOR”) functionality was validated by complementation experiments (Table 1, Fig. S3). Since these strains lack denitrification enzymes and are incapable of anaerobic growth, suspension assays were developed to culture bacteria aerobically while inducing gene expression prior to placement in non-growing, anoxic conditions. Strains were provided exogenous NO via the small molecule donor DETA NONOate (C_4_H_13_N_5_O_2_) at sub-toxic concentrations (Fig. S4) and then incubated under anoxic conditions for 24 hours at 37°C before the headspace was sampled; see Table 2 and Methods for more detail.

Under these conditions, iFhp displayed SP values (10.45 ± 2.17, *n*=5) that were significantly more positive than iNOR (-2.60 ± 5.41, *n*=5; *P*=0.004; Fig. 1E, top). iNOR values were also consistent with both our Δ*nosZ*Δ*fhp* measurements and prior *in vitro* NOR measurements (*14*). In addition, we observe a large variation (on the order of 10‰) in SP between biological replicates of NOR, in agreement with prior studies (-5 and -9‰; *n* = 2 in (*14*)). This variation neither correlates with the degree of nitrate consumption for Δ*nosZ*Δ*fhp*, nor N_2_O production for Δ*nosZ*Δ*fhp* and iNOR (Fig. S5), indicating that this variation in SP may be inherent to NOR.

Next, to validate Fhp SP values outside *Pa*, two WT, non-denitrifying strains with only Fhp, *Staphylococcus aureus* USA300 LAC and *Acinetobacter baumannii* ATCC 17978 were also measured. Fhp from *S. aureus* has 31.6% amino acid sequence similarity to Fhp from *P. aeruginosa*, while Fhp from *A. baumannii* has 98.5% similarity. However, all Fhps share a common catalytic site for NO binding and reduction, a globin module with heme B (*10*), that is responsible for imparting the observed SP. The SP of *S. aureus* (5.56 ± 7.21‰, *n*=3) and *A. baumannii* (10.38 ± 9.05‰, *n*=3) were both positive and statistically indistinguishable from *Pa* iFhp (Fig. 1E, bottom).

## Exogenous NO shifts SP values towards Fhp

Given the potential for Fhp to impart a positive SP distinct from NOR, we next sought to identify physiological conditions where it might dominate the N_2_O isotopic fingerprint in the WT. Historically, N_2_O isotopic measurements from pure cultures have been made for actively growing cells, which would be expected to amplify isotopic signatures imparted by catabolic enzymes like NOR. Yet evidence is mounting that slow, survival physiology dominates microbial existence in diverse habitats (*19, 20*), motivating N_2_O SP measurement during non-growth conditions.

To test if *Pa* can produce positive SP values indicative of Fhp activity, we grew WT *Pa* in denitrifying batch cultures and non-growing, anoxic suspensions with varying combinations of nitrate (NO_3_^-^) and DETA NONOate to provide NO endogenously via denitrification and/or exogenously via small molecule-mediated NO release (Fig. 2A), which we hypothesized would promote NOR or Fhp activity, respectively. We validated the induction of NOR and Fhp using quantitative unlabeled proteomics (Methods) and calculated the ratios of Fhp to NOR to quantify relative changes of each NO reductase. In denitrifying, batch culture conditions (Fig. 2B), the ratio of Fhp to NOR was less than one (∼0.25) and did not significantly change upon addition of NO (*P* = 0.09; Fig. 2B). By contrast, NorB, which contains the catalytic subunit of NOR, was undetectable before NO addition in the suspension assays (Fig. S6), which were performed by shifting oxic pre-grown cultures to non growing, anoxic conditions. Although NorB increased to detectable levels upon the addition of DETA NONOate (Fig. S6), Fhp was far more abundant, leading to a high ratio of Fhp to NOR (∼3, Fig. 2C).

**Fig. 2.**
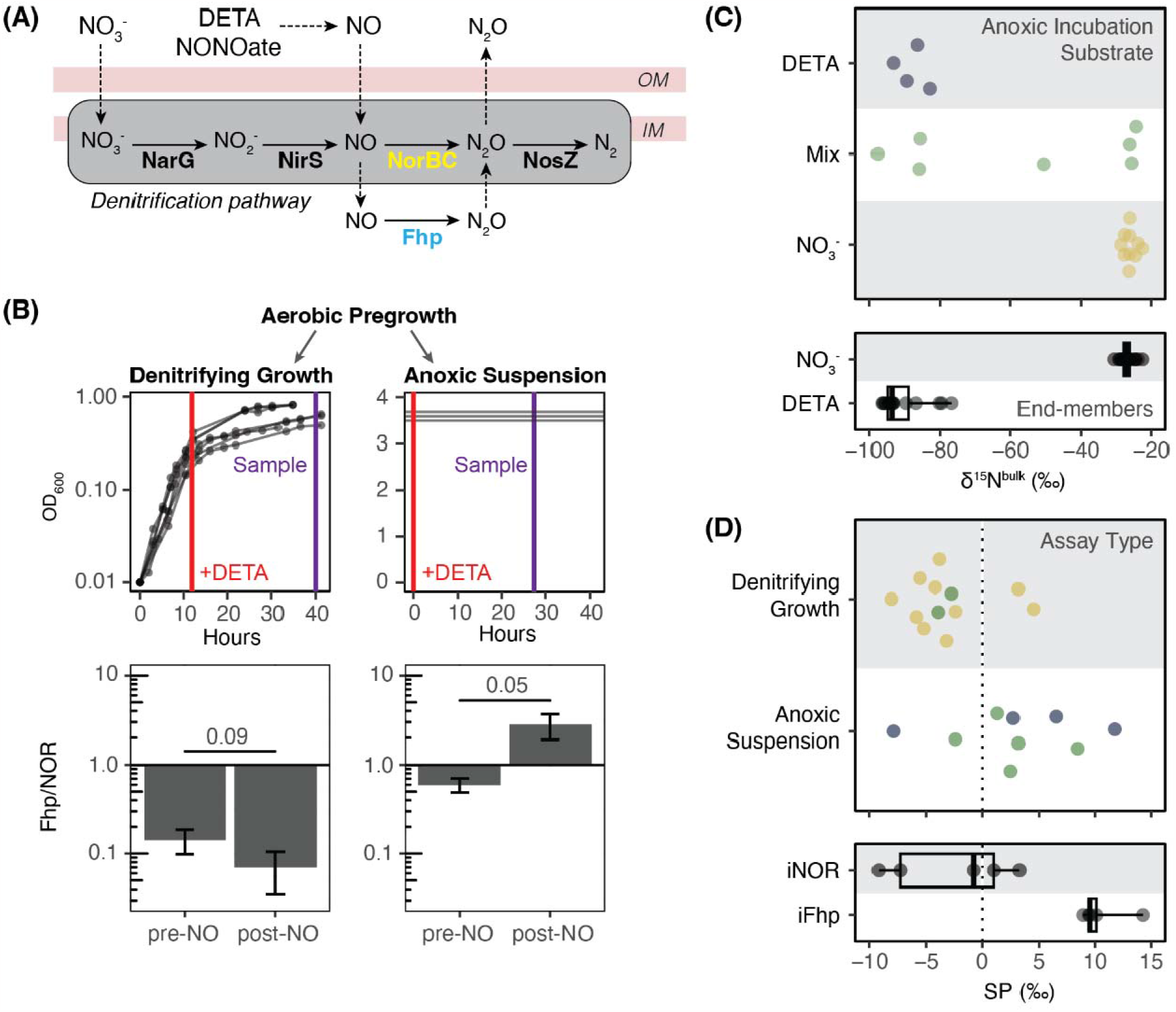
High concentrations of NO shift SP values towards Fhp. **(A)** In *Pa*, NorBC contributes to overall cell energetics as part of the denitrification pathway; Fhp does not and is primarily used for NO detoxification. **(B)** WT *Pa* was cultured anaerobically via two assay types after aerobic pre-growth with nitrate to either maximize growth via denitrification (left) or be re-suspended as non-growing cells (right). Exogenous NO was supplied through DETA NONOate (red lines) and headspace was then sampled for SP analysis (purple lines). Culture aliquots for proteomics analysis were taken immediately prior to NO addition (“pre-”) or during the same time as headspace sampling (“post-NO”). Ratio of Fhp to NOR in these conditions are shown as bar charts below; see Fig. S6 for full results. *P* values were calculated via Welch’s t-test. **(C)** δ^15^N^bulk^ values for WT *Pa* incubated anoxically with DETA (blue), nitrate (yellow) or both (green); end-member values are from non-WT *Pa* strains incubated with only nitrate or DETA as an NO source (Fig. S8). **(D)** SP measurements for WT *Pa* grown as denitrifying growths or anoxic suspensions, as illustrated in (B). Colors indicate anoxic incubation substrate and are the same as panel (C). iNOR and iFhp SP values are from Fig. 1E. For (C, D), box plots indicate median, upper and lower quartiles, and extreme values.

Paired SP and δ^15^N^bulk^ data allowed us to track which pool of NO was used by Fhp or NOR for N_2_O production (Fig. S7, Fig. 2C,D). When N-oxides are reduced to N_2_O, δ^15^N^bulk^ retains the isotopic signature of the original N(*21*)(*21*). The NO_3_^-^ and DETA NONOate used in our experiments had distinct δ^15^N values (0.40 ± 1.28 vs. -22.95 ± 0.15‰ respectively); non-WT *Pa* strains grown as batch cultures with only nitrate or incubated as suspension assays with DETA NONOate retained these distinct signatures in their δ^15^N^bulk^ values (-27.4 ± 1.4 vs. -91.0 ± 6.5‰ respectively, Fig. S8). All strains exhibit a large (roughly -70‰) of bulk δ^15^N when producing N_2_O from exogenous NO sourced from DETA NONOate, whether NOR or Fhp is utilized; this compares with typical bulk δ^15^N fractionations of -10 to -30 ‰ between source nitrate and product N_2_O in prior studies of biological nitrate reduction (*22*). Thus, the large amplitude contrast in δ^15^N between N_2_O produced from nitrate vs. exogeneous NO in our experiments reflects the combined influences of the difference in δ^15^N between our nitrate and NO substrates, and the difference in overall isotopic fractionations between the nitrate-to-N_2_O and the NO-to-N_2_O reactions. When WT *Pa* was incubated anoxically with either NO_3_^-^ or DETA NONOate, δ^15^N^bulk^ values correspondingly showed only one NO source (Fig. 2C); when given both substrates simultaneously, N_2_O could be made from varying ratios of both exogenous and endogenous NO.

SP data (Fig. 2D) was consistent with denitrifying batch cultures favoring NOR production, and non-growing, anoxic suspension assays favoring Fhp. When WT *Pa* was grown under denitrifying conditions, SP values were more negative and within the range of iNOR. However, in suspension assays, SP values spanned the range from iNOR to iFhp, consistent with increased Fhp abundance in these conditions. The most positive SP values, within the range of iFhp, were seen when WT *Pa* was given a high dose of both endogenous and exogenous NO in oxic pre-growth (NO_3_^-^ and DETA NONOate, blue stars, Fig. S7) followed by anoxic incubation with exogenous NO (DETA NONOate only, blue circles, Fig. 2D).

## Consequences for interpreting existing SP data

Because Fhp homologs are present in many denitrifying bacteria and AOB (Fig. S1, S9, Tables S1-S3), it is possible that Fhp may have contributed to the SP values measured in previous pure culture studies. Notably, all prior reports of SP from bacterial denitrifiers used strains that also have Fhp (Table S1); given the sensitivity of enzyme abundance to the physiological state during the time of measurement, it is plausible that the positive spread in SP values observed in these studies (*23*) may reflect cryptic Fhp activity. An Fhp homologue, Yhb, exists in yeast (*10*) and is present in previously studied fungal denitrifiers as well (Table S3), possibly contributing to the tail towards 10‰ observed from the literature (Fig. 1A) if the SP signature of Yhb is similar to that of Fhp.

Fhp is phylogenetically widespread and more abundant than NOR; therefore, measuring Fhp values from a representative group of diverse bacteria may illuminate the natural variation in SP values. In addition, measuring other NO-detoxifying proteins may shed further light on the SP values of this neglected class of non-catabolic enzymes. Flavo-diiron proteins, which only operate in anoxic conditions and only reduce NO to N_2_O for detoxification (*9*) present an attractive next target for SP measurements. Finally, further detailed studies of Fhp’s reaction mechanism paired with SP data is needed to reveal what determines the SP of N_2_O formation through NO reduction, for both abiotic and biotic reactions (*7, 24*–*26*).

Beyond helping to explain the N_2_O SP variation seen in prior pure-culture studies, our finding that Fhp produces an intermediate SP value that overlaps with many natural SP measurements, particularly those found in soil (median Fhp SP=9.5‰, median soil SP=10.9‰; Fig. S10). This begs the question: How can we distinguish Fhp-generated N_2_O from that produced by a mixture of other enzyme sources in complex environments such as soils or infected tissues? This is a difficult task. Though we can infer whether certain enzymes may be present and active based on knowledge of what regulates their expression, in order to predict whether they are active in any given sample, we need to know the conditions experienced by cells *in situ*. For example, our work indicates that Fhp dominates the SP fingerprint when cells grown under oxic conditions subsequently encounter a concentrated pulse of NO under anoxia. Intriguingly, pulses of NO and N_2_O have been detected after wetting of dryland soils (*27, 28*) nd opportunistic pathogens are thought to experience NO bursts from different cell types in the human immune system (*29*). Yet to speculate on whether such pulses may trigger Fhp activity, we would need to be able to track *in situ* NO and oxygen concentrations at the microscale. Ultimately, knowledge of the relative abundance of NO reductases present at the protein level in any given sample where N_2_O SP is measured, paired with knowledge of microscale environmental states, will aid in source attribution.

Though this manuscript focuses on N_2_O, the general approach presented here exemplifies how particular microbial metabolic pathways may be forensically distinguished by means of intramolecular isotopic biosignatures within their products. Going forward, we envision such an approach, when applied to other metabolites of interest, may help us better understand how microbial activities shape diverse habitats, from soils to animal hosts.

## Supporting information

Site Preference Full Data

## Acknowledgments

We thank Colette L. Kelly for valuable guidance and help with the scrambling correction; Nami Kitchen for assistance with IRMS measurements; and Nathan Hart at the Caltech Glass Shop for building the vacuum flasks. We thank the Dr. Tsui-Fen Chou and Baiyi Quan at the Caltech Proteome Exploration Laboratory for assistance with proteomics-based experiments.

## Funding

National Science Foundation Graduate Research Fellowship Program (RZW), Jane Coffin Childs Memorial Fund for Medical Research Fellowship (ZRL), National Institutes of Health grant R01 HL152190-03 (DKN, JML)

## Author contributions

Conceptualization: DKN, JME, RZW, ZRL

Methodology: RZW, ZRL, SAW

Investigation: RZW, ZRL, SAW, DKN

Visualization: RZW, ZRL, DKN, JME

Funding acquisition: DKN, JME, RZW, ZRL

Project administration: DKN

Supervision: DKN, JME

Writing – original draft: RZW, ZRL

Writing – review & editing: RZW, ZRL, DKN, JME

## Competing interests

Authors declare that they have no competing interests.

## Data and materials availability

All data are available in the main text or the supplementary materials.

## Supplementary Materials

Materials and Methods

Supplementary Text

Figs. S1 to S18

Tables S1 to S13

## Notes

### Competing Interest Statement

The authors have declared no competing interest.

