## Supplemental Methods, Figures and Tables for "Widespread detoxifying NO reductases impart a distinct isotopic fingerprint on N_2_O under anoxia"

**The PDF file includes:**

Materials and Methods

Supplementary Text

Figs. S1 to S18

Tables S1 to S13

### Materials and Methods

#### 1. Medium and nitric oxide donors

We amended synthetic cystic fibrosis medium (“Base SCFM”) [(*1*)](https://sciwheel.com/work/citation?ids=2777132&pre=&suf=&sa=0) with 20 mM sodium succinate and trace metals to increase cell and N_2_O yields (“SCFM Amended”). A 1000x solution of the trace metal stock (Trace element sol. SL-10; DSMZ) at a total volume of 1000 mL comprised: 1) 10.00 mL of HCl (25%; 7.7 M); 2) 1.50 g of FeCl_2_ x 4 H_2_O; 3) 70.00 mg of ZnCl_2_; 4) 100.00 mg of MnCl_2_ x 4 H_2_O; 5) 6.00 mg of H_3_BO_3_; 5) 190.00 mg of CoCl_2_ x 6 H_2_O; 6) 2.00 mg of CuCl_2_ x 2 H_2_O; 7) 24.00 mg of NiCl_2_ x 6 H_2_O; 8) 36.00 mg of Na_2_MoO_4_ x 2 H_2_O; 9) 990.00 mL of distilled water. All strains in this study were grown in SCFM-A media. The small molecule NO donor DETA NONOate (C_4_H_13_N_5_O_2_, #82120 Cayman Chemical Company) was used in certain experiments. It decays following first order kinetics in a pH-dependent manner to release two moles of NO per mole of DETA NONOate (half life of 20 hours at 37°C and pH 7.4).

#### 2. Strain generation

We measured the SP of N_2_O produced by five strains of *Pa*, and two wild-type strains of *Staphylococcus aureus* and *Acinetobacter baumannii* (Table 1).

*Pseudomonas aeruginosa* UCBPP-PA14 was the wild-type (WT) and parent strain of all genetic manipulations done in this study. Individual and combinatory mutants of *Pa* nitrate reductase (*ΔnarGHJI*; *PA14_13780-13830)*, nitrite reductase (*ΔnirS*; *PA14_06750*, nitric oxide reductase (*ΔnorBC*; *PA14_16810*, *PA14_16830*) and nitrous oxide reductase (*ΔnosZ*, *PA14_20200*) were generated previously [(*2*)](https://sciwheel.com/work/citation?ids=15334927&pre=&suf=&sa=0). *ΔnosZΔfhp* has the additional deletion of *fhp*, the flavohemoglobin protein / nitric oxide dioxygenase (*PA14_29640*). Clean deletions were done using allelic exchange as previously described [(*3*)](https://sciwheel.com/work/citation?ids=5942822&pre=&suf=&sa=0); briefly ~1 kb fragments surrounding the gene of interest were amplified by PCR and Gibson cloned into pMQ30 [(*4*)](https://sciwheel.com/work/citation?ids=30022&pre=&suf=&sa=0). Deletion constructs were introduced into *Pa* via triparental conjugation, and *E. coli* plasmid and helper strains were selected against on VBMM containing 50 µg/ml gentamicin [(*5*)](https://sciwheel.com/work/citation?ids=2068590&pre=&suf=&sa=0). Resulting Gent^R^ *Pa* cells were plated on 10% sucrose LB agar to isolate recombinants and screened via PCR. See Table S5 for primers used. Another strain, *ΔnosZΔnorBC*, was also used but it did not grow appreciably in the anaerobic, batch culture growth condition (Fig. S2); therefore its SP was not measured.

Two strains with inducible expression were created to increase N_2_O production amounts for isotopic measurement (Fig. S3). Strains with inducible *fhp* (‘iFhp,’ to denote *P. aeruginosa* *ΔnosZΔfhpΔnor* *att*::*mTn7*(GentR,*fhp*)) and *norBCD* (‘iNOR,’ to denote *P. aeruginosa ΔnarΔnirΔnorΔnosZΔfhp att*::*mTn7*(GentR,*norBCD*)) were generated by, first, amplifying *fhp* or *norCBD* from *P. aeruginosa* genomic DNA. See Table S5 for primers used. PCR products were ligated into plasmid the miniTn7 plasmid pJM220 [(*5*)](https://sciwheel.com/work/citation?ids=2068590&pre=&suf=&sa=0) via Gibson cloning [(*4*)](https://sciwheel.com/work/citation?ids=30022&pre=&suf=&sa=0) 3’ of the *rhaB* promoter for rhamnose-specific expression. Plasmids were delivered to *P. aeruginosa* via triparental conjugation with *Escherichia coli* SM10(λpir) and SM10(λpir) pTNS1 [(*5*)](https://sciwheel.com/work/citation?ids=2068590&pre=&suf=&sa=0), and exconjugants were selected on LB agar supplemented with chloramphenicol (10 µg/ml) and gentamicin (20 µg/ml) and verified by PCR.

In addition, we measured the SP of N_2_O produced by two wild-type, non-denitrifying bacteria with only *fhp/hmp* annotated in their genomes – *Staphylococcus aureus* USA300 LAC (putative flavohemoprotein *SAUSA300_0234*) and *Acinetobacter baumannii* ATCC 17978 (putative flavohemoprotein *A1S_3085*) (both kindly provided by Eric Skaar, Vanderbilt University Medical Center). Strains were first screened for N_2_O production in the presence of NO (See “N_2_O Screen” below; Table S2).

#### 3. Culturing conditions

iNOR, iFhp, and non-*Pseudomonas* were first screened for N_2_O production before scaling up the culturing process for isotopic measurement. All strains were first grown to a high density (OD_600_ ~ 3-4) from glycerol freezer stocks in aerobic pre-growths (25 mL SCFM-A, 250 rpm shaking for 16 hours at 37°C). Cells were then pelleted and fully re-suspended into 25 mL of fresh media in sealed, glass 18 x 150 mm Balch tubes. The headspace was then purged with N_2_ gas to establish anoxia, and 500 𝜇M DETA NONOate was added. Balch tubes were incubated statically for 24 hours at 37°C. The headspace was then sampled on the vacuum line and distilled to concentrate N_2_O and CO_2_ in a preliminary distillation (see below for further detail).

Next, all isotopic measurements were performed on strains grown in one of two ways: (i) suspension assays or (ii) batch culture (Fig. S12). All strains were grown in SCFM-A, but the NO source (KNO_3_^-^ vs. DETA NONOate) varied per experiment. iNOR, iFhp, *A. baumannii*, and *S. aureus* were only grown as suspension assays. *ΔnosZ* and *ΔnosZΔfhp* were only grown as batch cultures. WT *Pa* was grown as both suspension assays and batch cultures. All anoxic incubations were performed in custom vacuum sampling flasks (Fig. S13). Vacuum flasks could not be sterilized through autoclaving because the flask cracked under high pressures. Therefore, flasks were instead sterilized with 80% ethanol, then dried overnight at 56°C and exposed to UV light in a sterile, laminar-flow hood for 10 minutes.

For suspension assays, strains were first grown in shaking, aerobic pre-growths for 16 hours at 37°C (OD_600_ ~3-4) in 150 mL SCFM-A. The aerobic pre-growths for iNOR, iFhp, *A. baumannii*, and *S. aureus* were supplemented with 100 mM KNO_3_. For WT PA14 suspension assays, pre-growth was supplemented with 100 𝜇M DETA NONOate, 100 𝜇M DETA NONOate and 100 mM nitrate, or 100 mM nitrate. Next, cells transferred to 50 mL conical tubes, pelleted for 15 minutes at 23°C and 6,800 xg, and resuspended in 150 mL fresh SCFM-A. 500 𝜇M DETA NONOate was added to iNOR, iFhp, *A. baumannii*, and *S. aureus* experiments; iNor and iFhp was also supplemented with 305 𝜇M L-rhamnose monohydrate (C_6_H_12_O_5_ · H2O (Sigma-Aldrich R3875-25G) to promote rhamnose-inducible expression of *norBC* or *fhp*. For WT PA14 suspension assays, either 500 𝜇M DETA NONOate or 500 𝜇M DETA NONOate and 100 mM nitrate was added. Following suspension setup, vacuum flask headspace was purged with N_2_ gas to establish anoxia, and flasks were incubated statically for 24 hours at 37°C before headspace sampling. See Fig. S14 for more detail.

For batch culture assays, strains were first grown in aerobic pre-growths of 5 mL SCFM-A with 100 mM nitrate for 16 hours at 37°C, 250 rpm shaking (OD_600_ ~3-4). Cells were then diluted to OD_600_ = 0.01 in vacuum flasks with 150 mL of SCFM-A, For *ΔnosZ* and *ΔnosZΔfhp*, 100 mM of KNO_3_ was added. For WT *Pa*, either 100 mM KNO_3_ or 500 𝜇M DETA NONOate and 100 mM KNO_3_ was added. Vacuum flask headspace was purged with N_2_ gas to establish anoxia and incubated statically at 37°C. Flasks were sampled twice: first, approximately 12 hours at end-exponential growth, and, second, approximately 40 hours at end-stationary. One WT *Pa* batch culture experiment, where 500 𝜇M DETA NONOate and 100 mM KNO_3_ were added to the vacuum flask, was only sampled at ~40 hours after the DETA NONOate was added at ~12 hours. See Fig. S14 for more detail. Additional moles of nitrate were accidentally added in the Aug192021 batch for a final concentration of 233 mM nitrate (Table S7); however, no difference in SP was observed (Fig. 1D).

#### 4. Headspace sampling and N_2_O distillation

N_2_O was distilled from the headspace samples on an ultra-torr vacuum line prior to isotopic analysis (Fig. S14). First, the sample was expanded onto the left side of the line (Step 1); higher pressure samples were sampled by taking multiple aliquots while lower pressure samples were fully exposed to the line. Next, non-condensables (i.e. N_2_, Ar) were removed by passing the sample over a trap in liquid nitrogen (LN_2_, T2 in Fig. S14. Then, the sample was passed back and forth over the ascarite tube and the ethanol / dry ice slurry trap (T3, Fig. S14) to remove CO_2_ and H_2_O. Each pass lasted four minutes. The sample was isolated from the vacuum and the directionality of the sample flow was determined by either submerging T1 (clockwise flow) or T2 (counter-clockwise flow) in LN2. The ascarite tube was re-made roughly every six samples; it consists of a ≈10” length pyrex tube of ⅜” gauge containing sodium hydroxide (Ascarite II CO2 Absorbent, Thomas Scientific) and sealed with quartz wool on both ends. The ethanol / dry ice trap was a slurry of 100% ethanol (v/v) mixed with dry ice (solid CO_2_). Finally, the sample was passed over the ethanol / dry ice slurry for a final time and flame-sealed into a pyrex glass finger until isotopic analysis.

Two vacuum distillation blanks (0100, 0101) and a no-cells vacuum flask blank (0112) were measured to test if the distillation process causes significant isotopic fractionation (Table S8). A total mixture of 640 μmol CO_2_ and 290 μmol N_2_O were expanded to a total volume of 127 cc on the vacuum line, then equilibrated with a pyrex finger of ~5 cc containing room air and ~0.1 mL DI water. Two aliquots of this mixture were taken as mock samples (0100 and 0101). The no-cells vacuum flask blank (0112) was prepared as a batch culture, but after the headspace was purged with N_2_ gas, N_2_O from the reference tank was injected into the flask. This flask was then incubated at 37°C for ~12 hours and sampled at end-exponential phase (~12 hours). 0100 and 0101 showed little difference from the original N_2_O gas (roughly 0.1 ± 0.5‰ difference), indicating that the distillation process does not significantly fractionate our target gas. 0112 showed a -2.25 ± 0.90‰ difference in δ^18^O; this may have been caused by exchange of O isotopes between the incubated N_2_O gas and H_2_O – therefore our study relies on interpretation of the N isotopes instead.

#### 5. Site Preference measurements

#### 5.1 Delta (δ) Notation and Definition of Site Preference (SP)

All isotopic measurements in this study are reported in the delta notation (δ) in units of per mille (‰) where:

$\delta^{15}N=\left( \frac{{}^{15}R_{sam}}{{}^{15}R_{ref}}-1 \right)\times1000$

Equation S1

Where *^15^R* is the ratio of ^15^N/^14^N in the sample (“sam”) or reference (“ref”). All values here are reported to the international reference of Air for nitrogen.

Site Preference (δ^15^N^SP^ or “SP” in this study) is defined as the relative, intramolecular enrichment of the rare, stable isotope ^15^N for the central vs. terminal nitrogen in the linear, asymmetrical N_2_O molecule. To be consistent with prior work, we use the designations as defined by [(*6*, *7*)](https://sciwheel.com/work/citation?ids=1140081,1273457&pre=&pre=&suf=&suf=&sa=0,0) where the terminal nitrogen is labeled 𝜷, and the central nitrogen is labeled ⍺. In this convention, the ^15^R ratios for each site is defined as:

${}^{15}R^{\alpha}=\frac{[^{14}N^{15}N^{16}O]}{[^{14}N^{14}N^{16}O]}$

Equation S2

${}^{15}R^{\beta}=\frac{[^{15}N^{14}N^{16}O]}{[^{14}N^{14}N^{16}O]}$

Equation S3

Therefore, in delta notation:

$\delta^{15}N^{\alpha}=\left( \frac{{}^{15}R_{sam}^{\alpha}}{{}^{15}R_{ref}^{\alpha}}-1 \right)\times1000$

Equation S4

$\delta^{15}N^{\beta}=\left( \frac{{}^{15}R_{sam}^{\beta}}{{}^{15}R_{ref}^{\beta}}-1 \right)\times1000$

Equation S5

Site Preference is as defined as in [(*7*)](https://sciwheel.com/work/citation?ids=1273457&pre=&suf=&sa=0) :

$SP\equiv\delta^{15}N^{\alpha}-\delta^{15}N^{\beta}$

Equation S7

#### 5.2 Correction to international reference frame

The working reference gas (“Caltech Ref Gas”) used in this study was previously characterized relative to the international working standards for nitrogen and oxygen isotopes (Air and VSMOW respectively) by Tokyo Tech [(*8*, *9*)](https://sciwheel.com/work/citation?ids=5719748,1830225&pre=&pre=&suf=&suf=&sa=0,0). Values are listed in Table S9 below.

#### 5.3 SP Measurement Workflow

SP measurements were performed on two versions of the Thermo Scientific Ultra High-Resolution Isotope Ratio Mass Spectrometer (HR-IRMS), the ‘Prototype Ultra’ [(*10*)](https://sciwheel.com/work/citation?ids=5385218&pre=&suf=&sa=0) and the ‘Production Ultra.’ Two measurements were performed on each sample – the first at Mass 30 and 31 for δ^15^N^⍺^, and the second at Mass 44, 45 and 46 for δ^15^N^bulk^ and δ^18^O. All measurements were corrected for background (“Johnson”) noise [(*10*)](https://sciwheel.com/work/citation?ids=5385218&pre=&suf=&sa=0). Background correction was done before and after measurement on-peak to adjust for any pressure-related intensity changes, or other instrument changes over the course of the measurement. Gas injection pressures were calibrated to the main peak (Mass 30 or 44), and all sample measurements were bracketed by the reference gas.

A Mass 45 foot correction was done to correct for a ^13^C^16^O_2_ ‘foot’ that overlaps with the ^14^N^15^N^16^O / ^15^N^14^N^16^O measurement ‘shoulder.’ This ‘foot’ is present in both reference and sample gasses; reference gas was obtained through Matheson Gas at Ultra High Purity (99.99%). The correction was done by calculating a pressure-varying ratio of the foot vs. shoulder, then applying this pressure-varying ratio over the course of the measurement block. Two foot correction observations bracketing the measurement on-peak on the ‘shoulder’ were done to account for any pressure-related intensity changes over the course of the measurement. Over the observation period, both the foot and shoulder signal will decay exponentially with pressure, so both signals were fit with equations for exponential decay:

$I_{foot}=a_{foot}*EXP(b_{foot}*t)$

Equation S8

$I_{shoulder}=a_{shoulder}*EXP(b_{shoulder}*t)$

Equation S9

Where *I* = signal intensity, *t* = time, and *a* and *b* are fitted constants. We can then take the ratio of both equations for the correction:

$\frac{I_{foot}}{I_{shoulder}}=\frac{a_{foot}}{a_{shoulder}}*EXP[t*(b_{foot}-b_{shoulder})]$

Equation S10

Then this correction can be applied to the raw Mass 45 signal for the corrected Mass 45 signal:

$45_{corr}=\left[ 1-\frac{a_{foot}}{a_{shoulder}}*EXP[t*(b_{foot}-b_{shoulder})] \right]*45_{raw}$

Equation S11

This correction generally caused δ^15^N^bulk^ to become more negative by roughly 1‰.

#### 5.4 Shot noise error and limits of precision

In addition to background noise, shot noise is another inherent limit of isotope ratio measurements [(*10*–*12*)](https://sciwheel.com/work/citation?ids=14342148,14700453,5385218&pre=&pre=&pre=&suf=&suf=&suf=&sa=0,0,0). Therefore, for each measurement we calculated the shot noise error and compared it to the actual observed standard deviation of the measurement to see how close we approach shot noise limits. Figure S15 shows calculated shot noise vs. observed standard deviation for all measurements made for this study (samples, zero-enrichments, sample reruns; *n* = 79) across both Prototype and Production Ultras over six experimental sessions over two years. Linear regression across all points gives a slope close to 1 (*m* = 1.14 ± 0.04), indicating that overall measurements are approaching the shot noise error. Median calculated shot noise across all measurements (0.49‰, Fig. S15) is less than the median measured std. dev. (0.61‰, Fig. S15), indicating that, overall, measurements were reaching shot noise limits.

#### 5.5 Zero enrichment tests and instrument performance

‘Zero enrichment’ tests where the reference gas is measured as a sample against itself were regularly performed over the course of the study to ensure that pressure balance for the sample and reference gas bellows were correctly calibrated – i.e., if the bellows are correctly pressure calibrated, we would expect each measurement (δ^15^N^bulk^, δ^18^O and δ^15^N^ɑ^) to give a value of 0‰ within uncertainty (1 s.d.). Fig. S16 shows the result of eight zero enrichment tests for A) δ^15^N^bulk^, C) δ^18^O and E) δ^15^N^ɑ^ run on the Prototype (pink circles) and Production Ultra (blue circles). Results are largely 0‰ within uncertainty (1 s.d.) with the exception of δ^15^N^bulk^ and δ^18^O for Jun. 2021, and δ^15^N^bulk^ for Mar. 2023. However, this offset is on the order of 0.1‰, which is within the uncertainty of our shot noise error and is therefore likely due to the inherent limits of precision in our measurement. In addition, zero enrichments were run over a range of bellow pressures to gauge instrument performance across sample size (Fig. S16 B,D,F). A proxy for bellow pressure is ion beam intensity, since increased sample volume causes increased ion beam intensity. Calculating a linear correlation (not shown on figure) with minor ion intensity as the independent variable and δ values as the dependent variable gives adjusted R^2^ values of 0.09, 0.14 and -0.12 for δ^15^N^bulk^, δ^18^O and δ^15^N^ɑ^ respectively (calculation performed using R Statistical Software (v4.1.0; R Core Team 2021, [(*13*)](https://sciwheel.com/work/citation?ids=12719082&pre=&suf=&sa=0)) [call: *lm*()]. Therefore, there is no correlation between minor ion intensity and δ values, indicating that our measurement method is accurate across a range of sample sizes, bellow pressures, and signal intensities.

#### 5.6 Measurement consistency across instruments

Two samples, 0225 and 0230, were measured on both the Prototype and Production Ultras to gauge measurement consistency across instruments (Fig. S17). Samples were first measured in April 2022 on the Prototype Ultra. They were then removed from the sample bellow by freezing into a small glass finger with a finger-twist valve using liquid nitrogen (LN_2_). Samples were then taken to the vacuum line, frozen into glass break-seals using LN_2_, and stored in break-seals until further measurement. Samples were then re-measured on the Production Ultra on Dec. 2022. Measurements of 0225 and 0230 on both Ultras give the same value within measurement uncertainty (1 s.d.) vs. the reference gas for δ^15^N^bulk^, δ^18^O and δ^15^N^ɑ^, and all measurements approach the shot noise limit where std. dev. to shot noise ratio is 1 (Fig. S17). Standard deviation for both 0225 and 0230 is lower on the Prototype Ultra because there was more total sample in the first measurement (i.e. some sample was consumed during the first measurement).

#### 5.7 Scrambling Correction

Raw SP measurements must be corrected for ‘scrambling,’ a measurement artifact in which the ionization process in a gas source mass spectrometer “scrambles” all isotopologues of N_2_O, causing the inner (⍺) nitrogen and the outer (𝛽) nitrogen appear to be switched[(*14*)(*14*)](https://sciwheel.com/work/citation?ids=14439094&pre=&suf=&sa=0&dbf=0). There exist multiple strategies for scrambling corrections[(*15*–*18*)(*15*–*18*)](https://sciwheel.com/work/citation?ids=5247484,1026840,14308013,8346741&pre=&pre=&pre=&pre=&suf=&suf=&suf=&suf=&sa=0,0,0,0&dbf=0&dbf=0&dbf=0&dbf=0), and there are largely three levels of complexity that the correction can be performed at: i) A single-factor correction (𝛾) that assumes the scrambling behavior of ^14^N^15^N^16^O and ^15^N^14^N^16^O are equal, and that the contribution of ^17^O is negligible at Mass 31[(*7*, *19*)(*7*, *19*)](https://sciwheel.com/work/citation?ids=1273457,14897044&pre=&pre=&suf=&suf=&sa=0,0&dbf=0&dbf=0); ii) A two-factor correction (𝛾 and 𝜅) that accounts for the difference in scrambling between ^14^N^15^N^16^O and ^15^N^14^N^16^O, and assumes that ^17^O follows a mass-dependent relationship with ^18^O[(*20*, *21*)(*20*, *21*)](https://sciwheel.com/work/citation?ids=5714832,14743626&pre=&pre=&suf=&suf=&sa=0,0&dbf=0&dbf=0); and iii) A nine-factor correction that accounts for differences in scrambling between ^14^N^15^N^16^O, ^15^N^14^N^16^O, ^15^N^15^N^16^O, ^14^N^14^N^17^O, ^14^N^15^N^17^O, and ^15^N^14^N^17^O[(*18*)(*18*)](https://sciwheel.com/work/citation?ids=8346741&pre=&suf=&sa=0&dbf=0).

We used the single-factor correction following[(*6*, *7*)(*6*, *7*)](https://sciwheel.com/work/citation?ids=1273457,1140081&pre=&pre=&suf=&suf=&sa=0,0&dbf=0&dbf=0) because the nine-factor scrambling correction[(*18*)(*18*)](https://sciwheel.com/work/citation?ids=8346741&pre=&suf=&sa=0&dbf=0) requires measurement of up to nine external reference gasses, which we did not have, and because we believe the scrambling effects of ^15^N^15^N^16^O, ^14^N^14^N^17^O, ^14^N^15^N^17^O, and ^15^N^14^N^17^O are negligible at the level of precision needed for this study – i.e. the variations in SP between NOR and Fhp are on the order of 10‰. We did not use the two-factor correction following[(*20*, *21*)(*20*, *21*)](https://sciwheel.com/work/citation?ids=5714832,14743626&pre=&pre=&suf=&suf=&sa=0,0&dbf=0&dbf=0) because we were able to mass resolve ^17^O directly, and because that method is optimized for continuous-flow SP measurements.

We measured two replicates (RM5_1 and RM5_2) of external reference gas RM5[(*17*)(*17*)](https://sciwheel.com/work/citation?ids=14308013&pre=&suf=&sa=0&dbf=0) on the Production Ultra to calculate the scrambling factor (Table S10). Replicates were measured three months apart, and RM5_2 was measured at a lower sample amount. RM5 was used because it has a large, ~10‰ difference in δ^15^N between δ^15^N^bulk^ and δ^15^N^⍺^. We measured δ^15^N^bulk^ within uncertainty for RM5_2, and close to within uncertainty for RM5_1 (Table S10). We consistently measured the mean δ^15^N^⍺^ value to be more depleted by roughly 1‰, though all measured δ^15^N^⍺^ values overlapped with the reported δ^15^N^⍺^ value within uncertainty. This caused the SP values of RM5_1 and RM5_2 to be ‘compressed’ towards 0‰ compared to its reported value. δ^18^O values for RM5_1 and RM5_2 were measured to be their reported values within uncertainty (Table S10).

We then followed[(*7*)(*7*)](https://sciwheel.com/work/citation?ids=1273457&pre=&suf=&sa=0&dbf=0) to calculate 𝛾 = 0.04 ± 0.08 for RM5_1, and 𝛾 = 0.05 ± 0.11 for RM5_2. We therefore use an average 𝛾 value of 0.045 ± 0.136 for samples measured on the Production Ultra.[(*8*, *9*)(*8*, *9*)](https://sciwheel.com/work/citation?ids=5719748,1830225&pre=&pre=&suf=&suf=&sa=0,0&dbf=0&dbf=0) used the Prototype Ultra and measured samples in similar tuning conditions as used in this study; they used a one-factor correction of 0.110 ± 0.002. We therefore used 𝛾 = 0.110 ± 0.002 for samples measured on the Prototype Ultra. 𝛾 was likely lower on the Production Ultra because it has a lower baseline source pressure than the Prototype Ultra (3x10^-10^ vs. 9x10^-8^ mbar respectively).

The one-factor scrambling correction was performed as follows; an example correction is shown for one measurement of iFhp (Table S11). First, the sample is measured vs. Caltech Ref Gas. Next, values are corrected to international standards (AIR for N, VSMOW for O) using values reported by Tokyo Tech values (Table S11). Finally, the scrambling-adjusted ^15^R^⍺^ value (*^15^R^⍺^_adj_*) is calculated from the measured value (*^15^R^⍺^_meas_*) and the measured bulk value (*R^bulk^_meas_*):

$$R_{adj}^{\alpha}=\frac{R_{meas}^{\alpha}-2\gamma R_{meas}^{bulk}}{-2\gamma+1}$$

Equation S12

The final reported values, with scrambling correction and reported vs. AIR, are shown in the rightmost column of Table S11.

We checked our scrambling-corrected values against previously reported *in vitro* values for NOR. *ΔnosZΔfhp*, which only has NOR, was corrected using 𝛾 = 0.110 and iNOR was corrected using 𝛾 = 0.045. All corrected values overlap with previous *in vitro* measurements of a NOR enzyme purified from *Paracoccus denitrificans* ATCC 35512[(*22*)(*22*)](https://sciwheel.com/work/citation?ids=2285408&pre=&suf=&sa=0&dbf=0) (Fig. S18). In addition, as an internal check, the two samples measured on both the Production and Prototype Ultras (0225 and 0230, Fig. S17) gave similar values, implying that the scrambling factors are similar on both instruments. Indeed, 𝛾 = 0.045 ± 0.136 for the Production Ultra and 𝛾 = 0.110 ± 0.002 are similar within uncertainty.

#### 6. Isotopic measurement of DETA NONOate and nitrate substrates

We calculated the average δ^15^N of the initial NO reactant used in the suspension assays by measuring the difference in δ^15^N between the full and decomposed NO-donor, DETA NONOate (#82120, Cayman Chemical Company). DETA NONOate (C_4_H_13_N_5_O_2_) is a pH-dependent NO-donor that decays following first order kinetics to release two moles of NO per mole of DETA NONOate. At pH 7.4, it has a half life of 20 hours at 37°C and a half life of 56 hours at 22-25°C. pH was adjusted using NaOH and HCl, then the sample was prepared into 4x6 mm pressed tin capsules (Costech Analytical Technologies) for analysis. The δ^15^N of the full molecule, which has five Nitrogens, and the decayed molecule, which was three nitrogens, was measured on a Delta-V Advantage with Gas Bench and Costech elemental analyzer. Before measurement, the instrument was tuned with an internal standard to ensure instrument sensitivity and linearity, and to ensure correct measurement mass position. Three analytical replicates of each sample were measured. All samples were bracketed at the beginning and end of the run by a suite of external isotope standards (Urea δ^15^N = 0.0‰; Acetanilide δ^15^N = 19.56 ± 0.03‰; all reported vs AIR), tin capsule blanks, and NaOH and HCl blanks. After correcting for blanks, measured δ^15^N values were then corrected to reported values vs. AIR using the external Urea and Acetanilide standards. On average, the correction decreased the measured δ^15^N values by 0.2‰.

We then calculated the average δ^15^N of the released NO molecules by mass balance using the equation:

$2*\delta^{15}N_{2N}=5*\delta^{15}N_{5N}-3*\delta^{15}N_{3N}$

Equation S13

Where δ^15^N_2N_, δ^15^N_5N_, and δ^15^N_3N_ refer to the average δ^15^N of the released NO molecules, the full DETA NONOate molecule, and the decayed DETA NONOate molecule respectively. All results are reported in Table S12. Nitrate δ^15^N values were measured in the same way, except pH was not adjusted (Table S7).

Non-WT *Pa* strains were incubated with only nitrate or DETA NONOate as the NO source to determine if the δ^15^N signal of the NO substrate was inherited in δ^15^N^bulk^ (Fig. S8). *A. baumannii*, iNOR, iFhp and *S. aureus* were all grown as suspension assays and showed δ^15^N^bulk^ values of -91.0 ± 6.5‰ (mean ± s.d.). Though strains were incubated anoxically with both DETA NONOate and nitrate, NO was only sourced from DETA NONOate because nitrate could not be reduced to NO – i.e. native denitrification pathway was deleted in iNOR and iFhp, and does not exist in wild-type *A. baumannii* and *S. aureus* strains. 𝛥*nosZ* and 𝛥*nosZ*𝛥*fhp* strains were grown as batch cultures and incubated anoxically with only nitrate and showed δ^15^N^bulk^ values of -27.4 ± 1.4‰. This implies a δ^15^N fractionation factor of -27.8 ± 1.90‰ for nitrate to N_2_O, and -68.1 ± 6.5‰ for DETA NONOate to N_2_O.

#### 7. Rayleigh plots of NOR-only strains

Rayleigh plots of two strains with only NOR, Δ*nosZ*Δ*fhp* and iNOR, were made to test if the variation in SP followed a Rayleigh distillation relationship [(*23*)](https://sciwheel.com/work/citation?ids=8975346&pre=&suf=&sa=0). Δ*nosZ*Δ*fhp* was grown in batch culture denitrifying conditions while iNOR was grown as a suspension assay.

For Δ*nosZ*Δ*fhp*, the fraction of nitrate consumed was calculated two ways – from the total amount of N in each aliquot as measured on the Delta-V Advantage (*f_nitrate_*), or from the moles of N_2_O distilled from the headspace as measured in the direct-injection bellows on the Prototype or Production Ultra (*f_N2O_*). After each sampling time point for Δ*nosZ*Δ*fhp* where N_2_O measurement was performed (end-exponential and end-stationary), a roughly 5 mL aliquot of the liquid culture was taken and immediately flash frozen in liquid nitrogen. Aliquots were then kept frozen at -80°C until isotopic analysis. In addition, for each sampling batch, an aliquot KNO_3_, SCFM-A media, and DI water were flash frozen and stored as well. When samples were ready for analysis on the Delta-V Advantage with Gas Bench and Costech elemental analyzer as described above, all flash frozen aliquots were thawed at room temperature and aliquots were pipetted in triplicate into individual 5x9 mm pressed tin capsules (Costech Analytical Technologies) and left to dry overnight. Similar to above, raw measurements were corrected for tin capsule blanks using Urea and Acetanilide standards. Moles of total nitrogen were calculated based on the total peak area of each sample. This analysis cannot distinguish between N sourced from KNO_3_ or SCFM-A, though there are roughly four times more moles of N from KNO_3_ vs. SCFM-A (Table S7). The fraction of total N remaining was calculated by dividing the total N measured by the total N (KNO_3_ and SCFM-A) initially added. This is referred to as *f_nitrate_*.

We then calculated *f_N2O_* for Δ*nosZ*Δ*fhp*:

$f_{N2O}=1-\frac{2*n_{N2O}}{n_{NO3-}}$

Equation S14a

Where *n_N2O_* are the moles of N_2_O produced and *n_NO3-_* are the moles of nitrate initially added. N_2_O pressure was recorded in the sample bellows of the Ultra IRMS before each analysis, and moles of gas were calculated using the ideal gas law. Following similar studies, this equation assumes that every mole of nitrate that is taken up by the denitrification pathway results in two moles of N_2_O [(*24*)](https://sciwheel.com/work/citation?ids=13028172&pre=&suf=&sa=0).

*f_N2O_* was calculated in a similar manner for iNOR, except moles of N_2_O produced were compared to moles of DETA NONOate (*n_DETA_*) added.

$f_{N2O}=1-\frac{n_{N2O}}{{2*n}_{DETA}}$

Equation S14b

We then constructed SP Rayleigh curves [(*23*)](https://sciwheel.com/work/citation?ids=8975346&pre=&suf=&sa=0) following [(*24*, *25*)](https://sciwheel.com/work/citation?ids=13028172,1437434&pre=&pre=&suf=&suf=&sa=0,0), where SP is plotted against -(f*lnf)/(1-f), where *f* is the fraction of the remaining substrate. For Δ*nosZ*Δ*fhp*, two plots were made using the two different ways that *f* was calculated (*f_nitrate_* or *f_N2O_*). For iNOR, plot was only made using *f_N2O_* (Fig. S5).

Fitted values for *m* and *b* gave very large uncertainties and were of low confidence (Fig. S5). In particular, fitted values using *f_N2O_* had extremely large uncertainties due to the narrow range of the x-axis – i.e. nitrate was given at saturating conditions so the nitrate pool was not very depleted, resulting in *f_N2O_* values around 0.99. *f_nitrate_* varied over a larger range, likely because not all the nitrate consumed ended up as N_2_O (i.e. due to assimilatory nitrate processes, or from loss along the denitrification pathway), so the approach for calculating *f* in Eqn. S14a likely gives an overestimate. However, overall no correlation of SP with *f* is seen in the NOR-only strains, 𝛥nosZ𝛥*fhp* and iNOR.

#### 8. AnnoTree Search Parameters

A phylogram of species with annotated Fhp/Hmp sequences was first made from the NCBI database (Fig. S1, Table S13). Phylogram was made to include representative strains from a range of known bacterial species. The amino acid sequence of Fhp from *P. aeruginosa* PA14 was used (*PA14_29640*). Default NCBI protein BLAST blastp parameters were used to identify Fhp orthologs. Protein sequences were collected, and a simple phylogeny was constructed using EMBL-EBI Simple Phylogeny tool with default parameters and neighbor-joining clustering [(*26*)](https://sciwheel.com/work/citation?ids=12816401&pre=&suf=&sa=0). Two strains with a high and low amino acid sequence similarity were selected for further N_2_O screening and SP measurement. Fhp from *S. aureus* shows 31.6% sequence similarity to Fhp from *P. aeruginosa*, while Fhp from *A. baumannii* shows 98.5% similarity. Fhp and NorBC were also queried from AnnoTree, a functionally annotated database of >27,000 bacterial and >1,500 archaeal genomes [(*27*)](https://sciwheel.com/work/citation?ids=6807209&pre=&suf=&sa=0). Since Fhp from *S. aureus* shows 31.6% sequence similarity to Fhp from *P. aeruginosa*, the default search parameters were used: % identity: 30; E value: 0.00001; % subject alignment: 70; % query alignment: 70. Results are shown in Tables S1-S3 at the phylum level.

#### 9. Proteomics

Cells were collected in 5 mL aliquots from batch denitrifying or anoxic suspension assays immediately prior to DETA-NONOate addition (Fig S4, red line) or after 24-28 h incubation with DETA-NONOATE (Fig. S12, purple line), centrifuged at 6,800 xg, and pellets were frozen at -80°C. Thawed pellets were processed and digested via S-Trap^TM^ (ProtoFi, LLC, Fairport, NY) micro protocol digestion. Pellets were resuspended in lysis buffer (5% SDS, 25 mM TEAB pH 8.5) and sonicated for lysis. MgCl_2_ was added to (2 mM final) concentration and incubated at room temperature for 5 min. Samples were centrifuged for 10 min at 13,000 xg. Samples were then reduced with TCEP (5mM final), and samples were alkylated by addition of MMTS (20 mM final). Samples were acidified with phosphoric acid (2.5% final concentration), mixed with binding/wash buffer (100 mM TEAB, 90% methanol), and applied to the S-Trap column. Samples were centrifuged at 4,000 xg for 30 s and washed 3 times with binding/wash buffer. S-Trap was centrifuged for 1 min at 4,000 xg to fully remove binding/wash buffer. S-Trap was transferred to 1.7 mL microcentrifuge tube. Digestion buffer (1 µg trypsin/10 µg sample weight in 50 mM TEAB) was added to the S-Trap, and the tubes were loosely capped and incubated for 1 h at 47°C. 40 ul of elution buffer 1 (50 mM TEAB in water) was applied to S-Trap and centrifuged for 1 min at 4,000 xg. Elution and centrifugation was repeated with elution buffer 2 (0.2% formic acid in water), and elution buffer 3 (50% acetonitrile in water). Eluted peptides were pooled, dried, and resuspended in 0.2% formic acid for analysis.

LC–MS analysis of digested peptides was performed on an EASY-nLC 1200 (Thermo Fisher Scientific, San Jose, CA) coupled to a Q Exactive HF Orbitrap mass spectrometer (Thermo Fisher Scientific, Bremen, Germany) equipped with a Nanospray Flex ion source: 500 ng peptides of each sample were directly loaded onto an Aurora 25 cm × 75 μm ID, 1.6 μm C18 column (Ion Opticks) heated to 50°C. The peptides were separated with a 2 h gradient at a flow rate of 350 nl/min as follows: 2%–6% Solvent B (7.5 min), 6%–25% B (82.5 min), 25%–40% B (30 min), 40%–98% B (1 min), and held at 98% B (12 min). Solvent A consisted of 97.8% H_2_O, 2% ACN, and 0.2% formic acid and solvent B consisted of 19.8% H_2_O, 80% ACN, and 0.2% formic acid. The Q Exactive HF was operated in data-dependent mode with Tune (version 2.8 SP1 build 2806) instrument control software. Spray voltage was set to 1.6 kV, S-lens RF level at 50, and heated capillary at 275°C. Full scan resolution was set to 60,000 at *m*/*z* 200. Automatic gain control target was 3 × 10^6^ with a maximum injection time of 15 ms. Mass range was set to 375–1500 *m*/*z* and charge state inclusion set to select precursors of charge state 2–5 for DDA analysis. For data-dependent ms2 scans, the loop count was 12, AGC target was set at 1 × 10^5^, intensity threshold was kept at 1 × 10^5^, and dynamic exclusion set to exclude precursors after one time for 45 seconds. Isolation width was set at 1.2 *m*/*z* and a fixed first mass of 100 was used. Normalized collision energy was set at 28. Peptide match was set to off, and isotope exclusion was on. Data acquisition was controlled by Xcalibur (4.0.27.42), with ms1 data acquisition in profile mode and ms2 data acquisition in centroid mode.

Data analysis was performed using Thermo Proteome Discoverer 2.5 (Thermo Fisher Scientific, San Jose, CA) with a SEQUEST algorithm (PMID 24226387). The data was searched against the Pseudomonas aeruginosa UCBPP-PA14 proteome (UP000002438) acquired from UniProtKB (PMID: 36408920) in 2022-2-09. The ms1 matching tolerance was 20 ppm, and the ms2 tolerance was 0.02 Da. Carbamidomethyl (+57.021 Da) on cysteine was set as static modification, and Oxidation (+15.995 Da) on methionine was set as dynamic modification. Acetylation (+42.011 Da), Met-loss (-131.040 Da), and Met-loss + Acetylation (-89.030 Da) at protein N-terminus were also set as dynamic modifications. A maximum of 2 miss-cleavages were allowed in the search. A concatenated target decoy-based percolator was utilized to control the false discovery rate. The q-value cutoff was set as 0.05. Protein abundances were reported using ms1 feature-based label-free quantitation. The median abundance for each sample was normalized to the same value.

SUPPLEMENTARY FIGURES

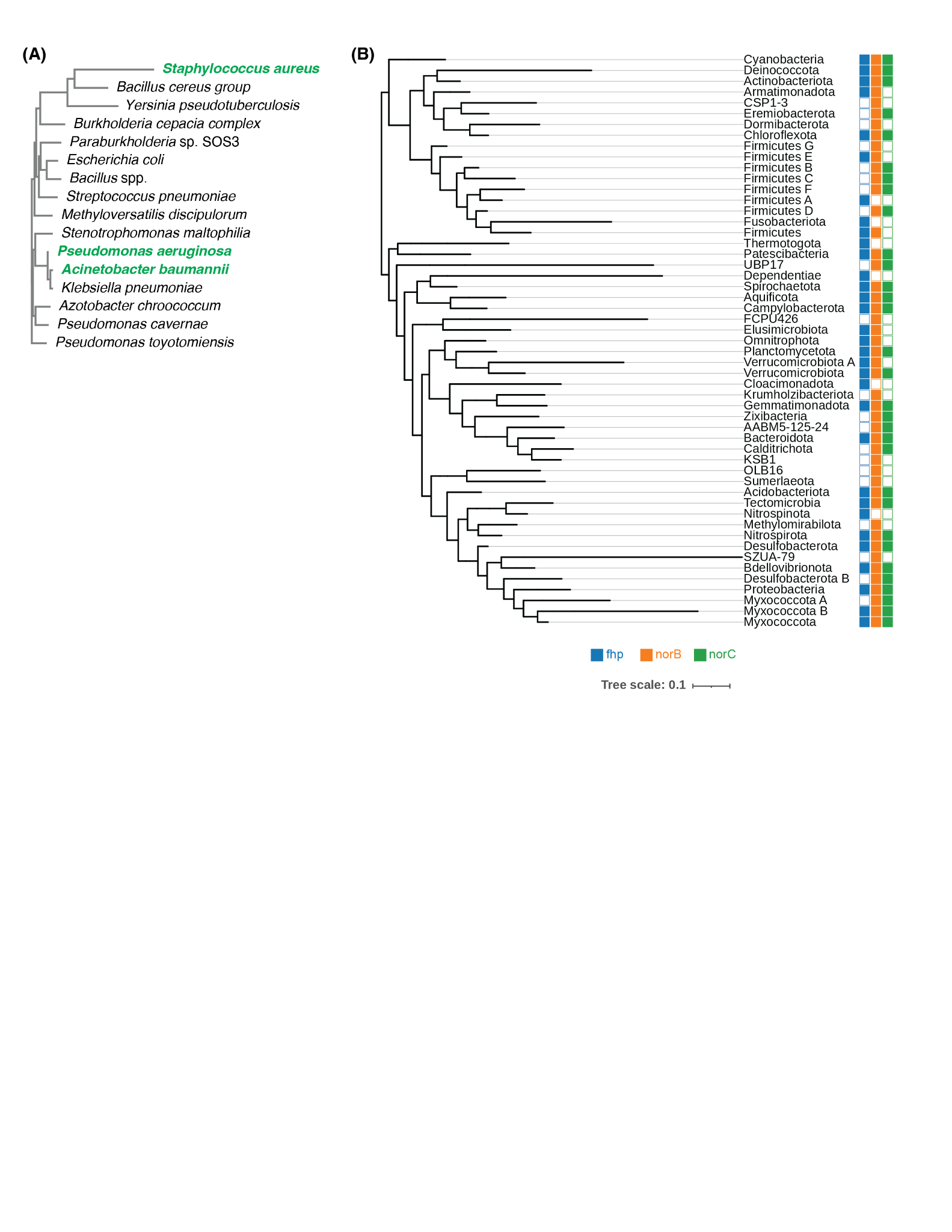

##### Fig. S1. Phylogeny of Fhp in Bacteria.

**(A)** Phylogram of annotated Fhp/Hmp amino acid sequences in the NCBI database; phylogram was curated to show a representative group of bacteria. Strains in green were measured for SP in this study. **(B)** Tree showing abundance of Fhp, NorB and NorC across Bacteria at the Phylum level, annotated in AnnoTree [(*27*)](https://sciwheel.com/work/citation?ids=6807209&pre=&suf=&sa=0) and visualized using the interactive tree of life (iTOL). Search parameters for AnnoTree were: % identity: 30; E value: 0.00001; % subject alignment: 70; % query alignment: 70.

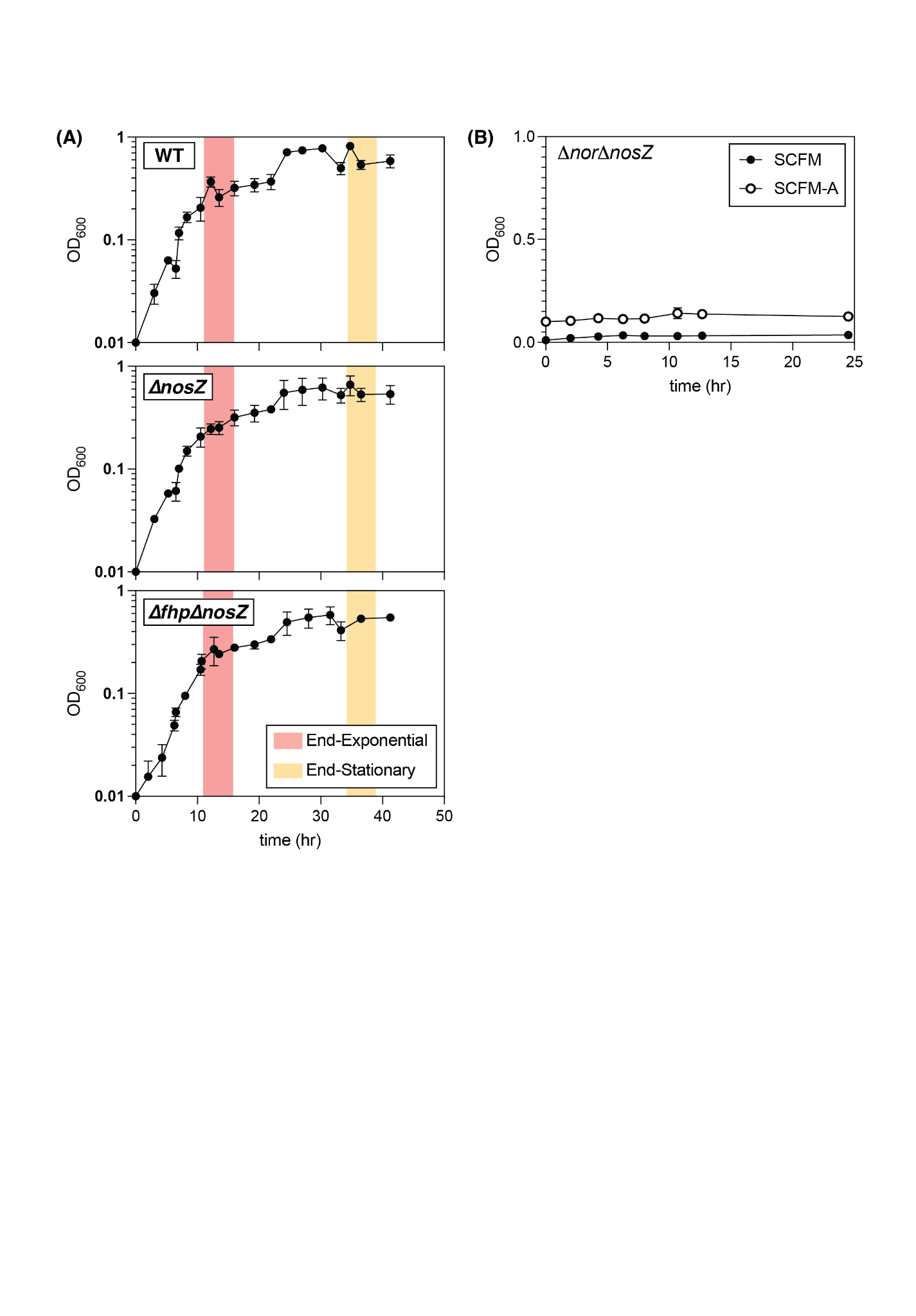

##### Fig. S2. Growth Curves.

**(A)** Growth curves measured by OD600 for WT *Pa*, 𝛥*nosZ*, and 𝛥*fhp*𝛥*nosZ* grown in batch culture, denitrifying conditions with headspace sampling times for SP measurements (end-exponential, red; end-stationary, yellow). Data points are mean +/- standard deviation (n=6). **(B)** Growth curves *𝛥nor𝛥nosZ* strain in SCFM (black) and SCFM-A (white) media. Data points are mean +/- standard deviation (n=3).

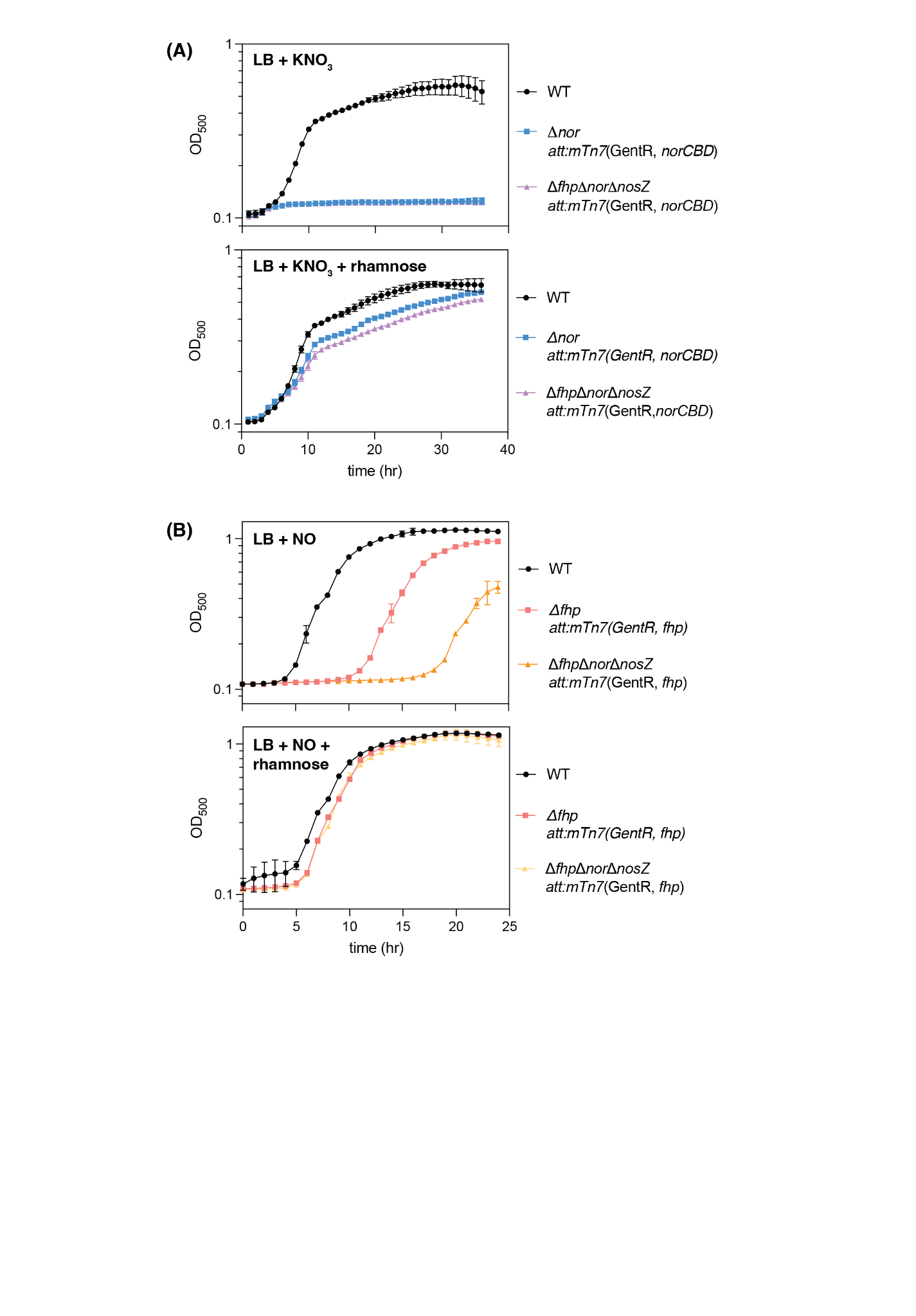

##### Fig. S3. Construction of iNOR and iFhp strains.

**(A)** Growth curve measured by OD500 of WT *Pa* (black), 𝛥*nor* chromosomally complemented with *nor* under a rhamnose-inducible promoter (*att*:*mTn7*(GentR, *norCBD*); blue) and 𝛥*fhp*𝛥*nor*𝛥*nosZ* (purple) with rhamnose-inducible *nor* grown anaerobically in Lysogeny Broth (LB) media with 40 mM potassium nitrate alone (upper panel) or 40 mM nitrate and 305 𝜇M rhamnose (lower panel). Data points are mean +/- standard deviation (n=3). **(B)** WT *Pa* (black), 𝛥*fhp* chromosomally complemented with *fhp* under a rhamnose-inducible promoter (*att*:*mTn7*(GentR, *fhp*); pink) and 𝛥*fhp*𝛥*nor*𝛥*nosZ* (yellow) with rhamnose-inducible *fhp* were grown aerobically in LB media with 500 𝜇M DETA NONOate alone (upper panel) or 500 𝜇M DETA NONOate and 305 𝜇M rhamnose (lower panel). Data points are mean +/- standard deviation (n=4).

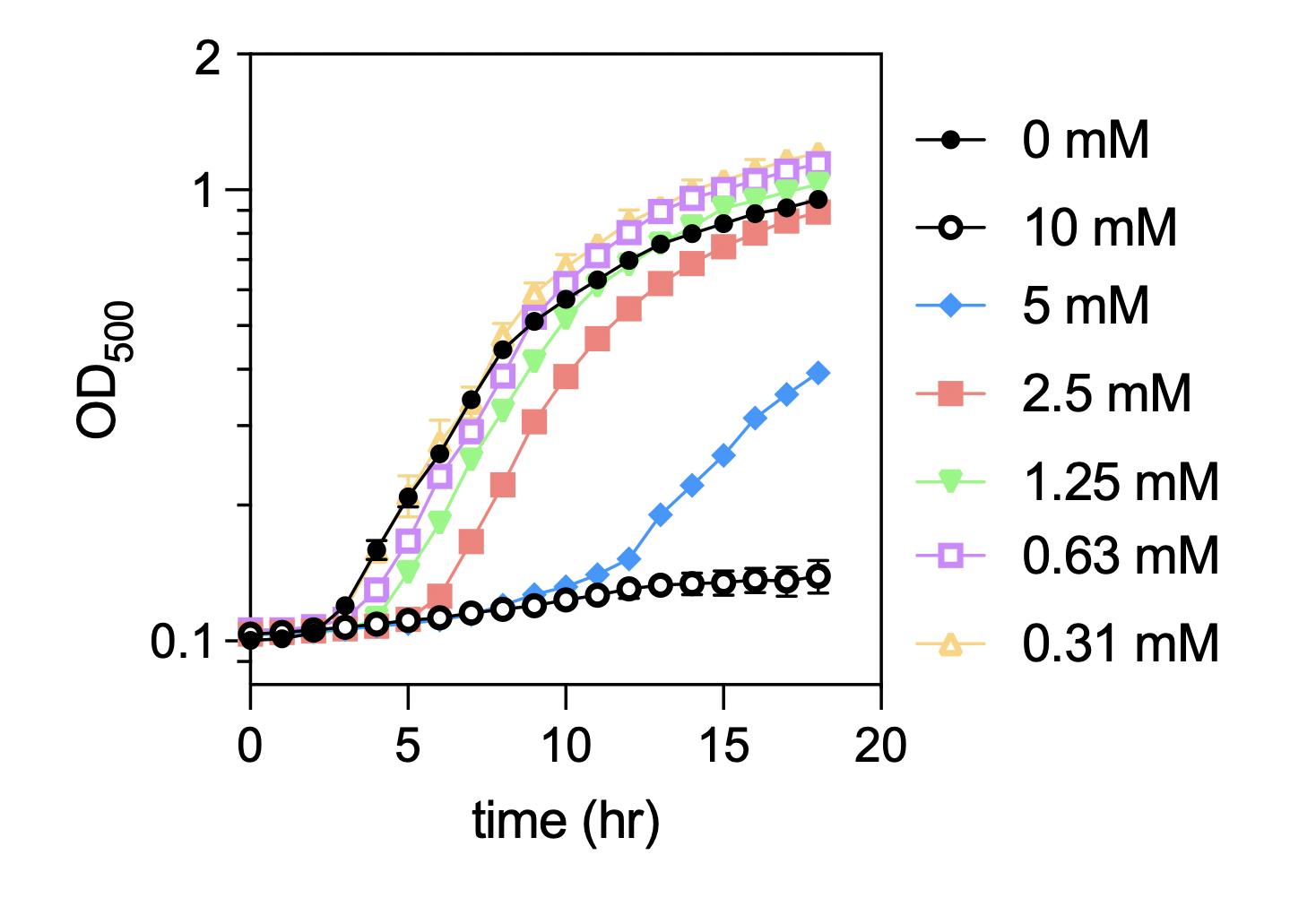

##### Fig. S4: Growth of WT *Pa* with DETA-NONOate titration.

WT *Pa* was grown in LB in the presence of DETA-NONOate with 2-fold dilutions ranging from ~300 𝜇M to 10 mM and OD500 monitored over time. A concentration of less than 1 mM DETA-NONOate did not appreciably affect growth.

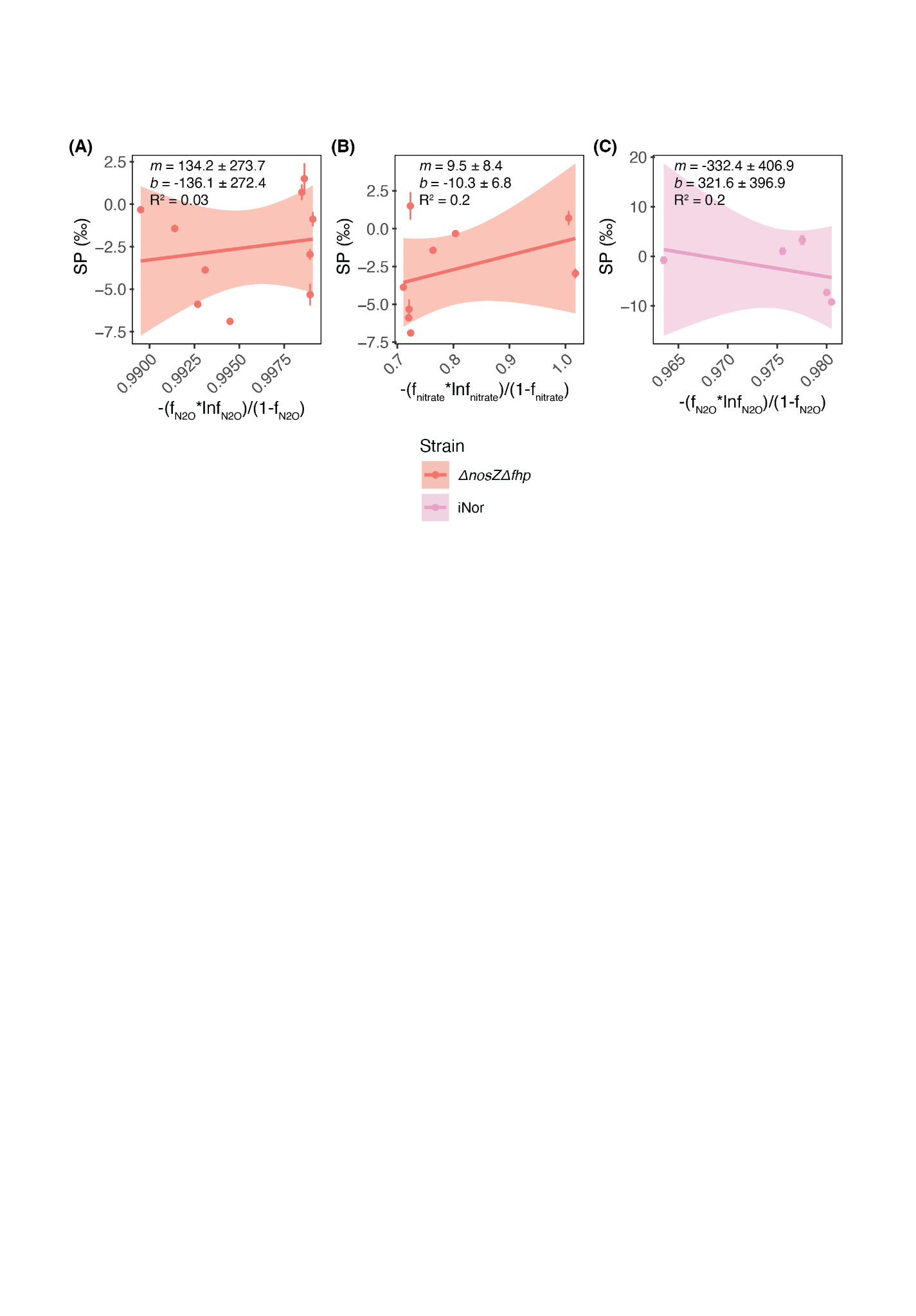

##### Fig. S5. Rayleigh plots of NOR-only strains.

Rayleigh plots for strains with only NOR: Δ*nosZ*Δ*fhp* **(A,B)** and iNOR **(C)**. *f* (fraction of substrate remaining) was calculated either from the amount of nitrate remaining in batch culture (*f_nitrate_*) or based on moles of N_2_O produced compared to the amount of nitrate or DETA NONOate initially added (*f_N2O_*_,_ Equations S14a,b). Results of linear regression are shown in the upper right corner of each plot; all analyses and data visualization were performed using R Statistical Software (v4.1.0; R Core Team 2021, [(*13*)](https://sciwheel.com/work/citation?ids=12719082&pre=&suf=&sa=0)) and the ggplot2 package (v3.3.6; Wickham, 2016, [(*28*)](https://sciwheel.com/work/citation?ids=14398114&pre=&suf=&sa=0)).

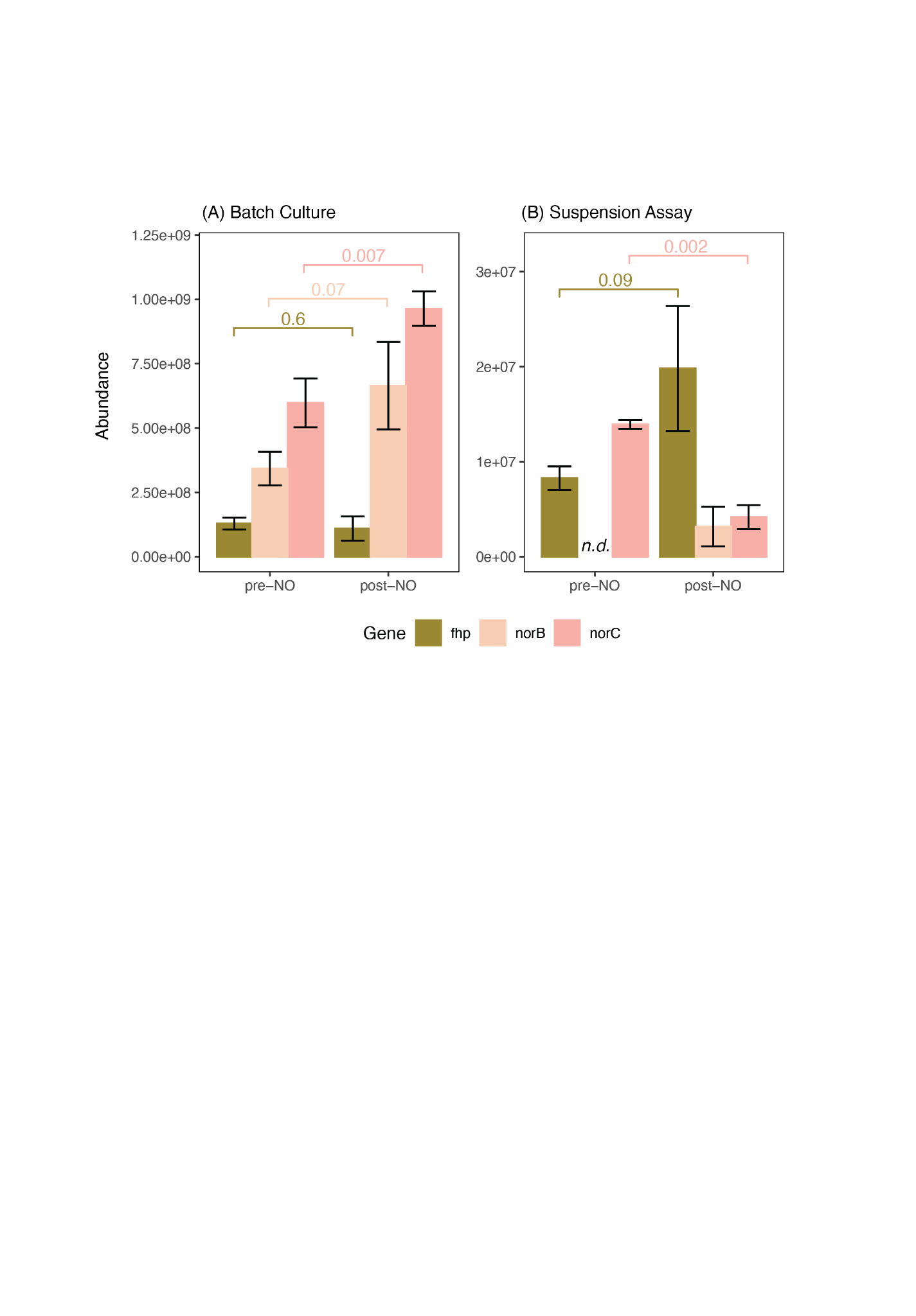

##### Fig. S6: Fhp, NorB and NorC protein abundances.

**(A)** Protein abundances for *fhp*, *norB* and *norC* before and after NO-addition for WT PA14 grown in batch culture. The ratio of *fhp* to *norB* and *norC* is presented in the main text. *P* values were calculated using Welch’s t-test for two independent groups and variance was not assumed to be the same across groups. See main text Fig. 2B for experimental set-up. **(B)** Results for WT PA14 grown in suspension assays; see main text Fig. 2C for experimental set-up. *P* value could not be calculated for *norB* in the suspension assay because it was not detected in any replicate; detection limit was 7264, 8974, and 15437 counts for the 3 replicates, respectively. For both panels, values represent the mean ± s.d. of three biological replicates. All analyses and data visualization were performed using R Statistical Software (v4.1.0; R Core Team 2021, [(*13*)](https://sciwheel.com/work/citation?ids=12719082&pre=&suf=&sa=0)) and the ggplot2 package (v3.3.6; Wickham, 2016, [(*28*)](https://sciwheel.com/work/citation?ids=14398114&pre=&suf=&sa=0)).

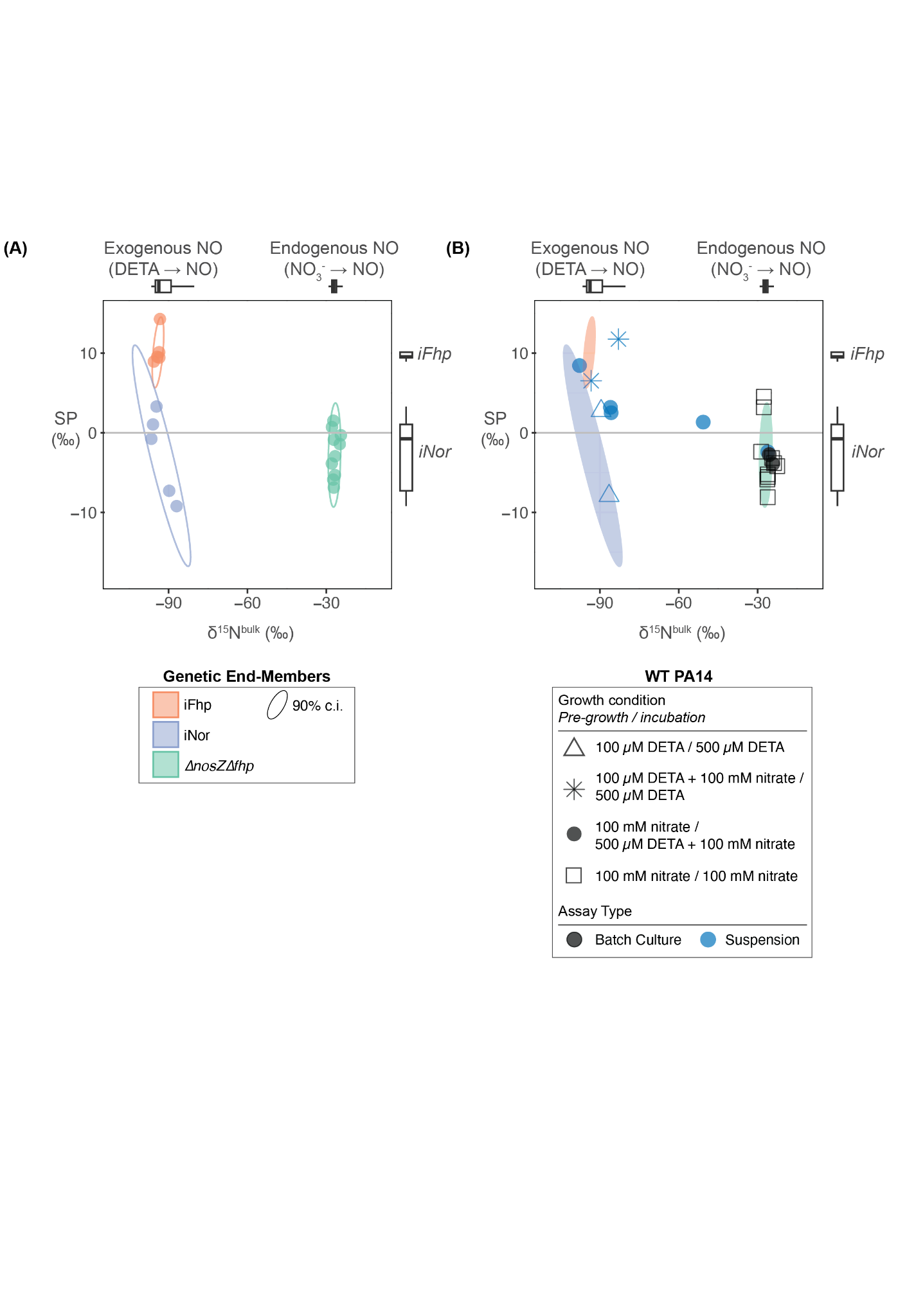

##### Fig S7: Paired SP and δ^15^N^bulk^ data for all experiments in parent strain *P. aeruginosa*.

**(A)** δ^15^N^bulk^ and SP data for all genetic end-member; i.e. strains with only NOR or Fhp (iFhp, iNor and 𝛥*nosZ*𝛥*fhp*). Ellipses show 90% confidence intervals (c.i.). Boxplots showing expected values for exogenous vs. endogenous NO source (above plot) are taken from Fig. S8, and boxplots showing iNor vs. iFhp values are taken from Fig. 1E. iFhp and iNor were grown as suspension assays while 𝛥*nosZ*𝛥*fhp* was grown as batch culture. **(B)** Overlay of 90% c.i. from (A) with experimental results from WT PA14, which has both NOR and Fhp. WT PA14 was grown as either batch culture or suspension assays (black vs. blue data points) with varying combinations of NO sources in the aerobic pre-growth vs. anoxic incubation (triangles, stars, circles, and squares as noted in the legend). All analyses and data visualization were performed using R Statistical Software (v4.1.0; R Core Team 2021, [(*13*)](https://sciwheel.com/work/citation?ids=12719082&pre=&suf=&sa=0)) and the ggplot2 package (v3.3.6; Wickham, 2016, [(*28*)](https://sciwheel.com/work/citation?ids=14398114&pre=&suf=&sa=0)).

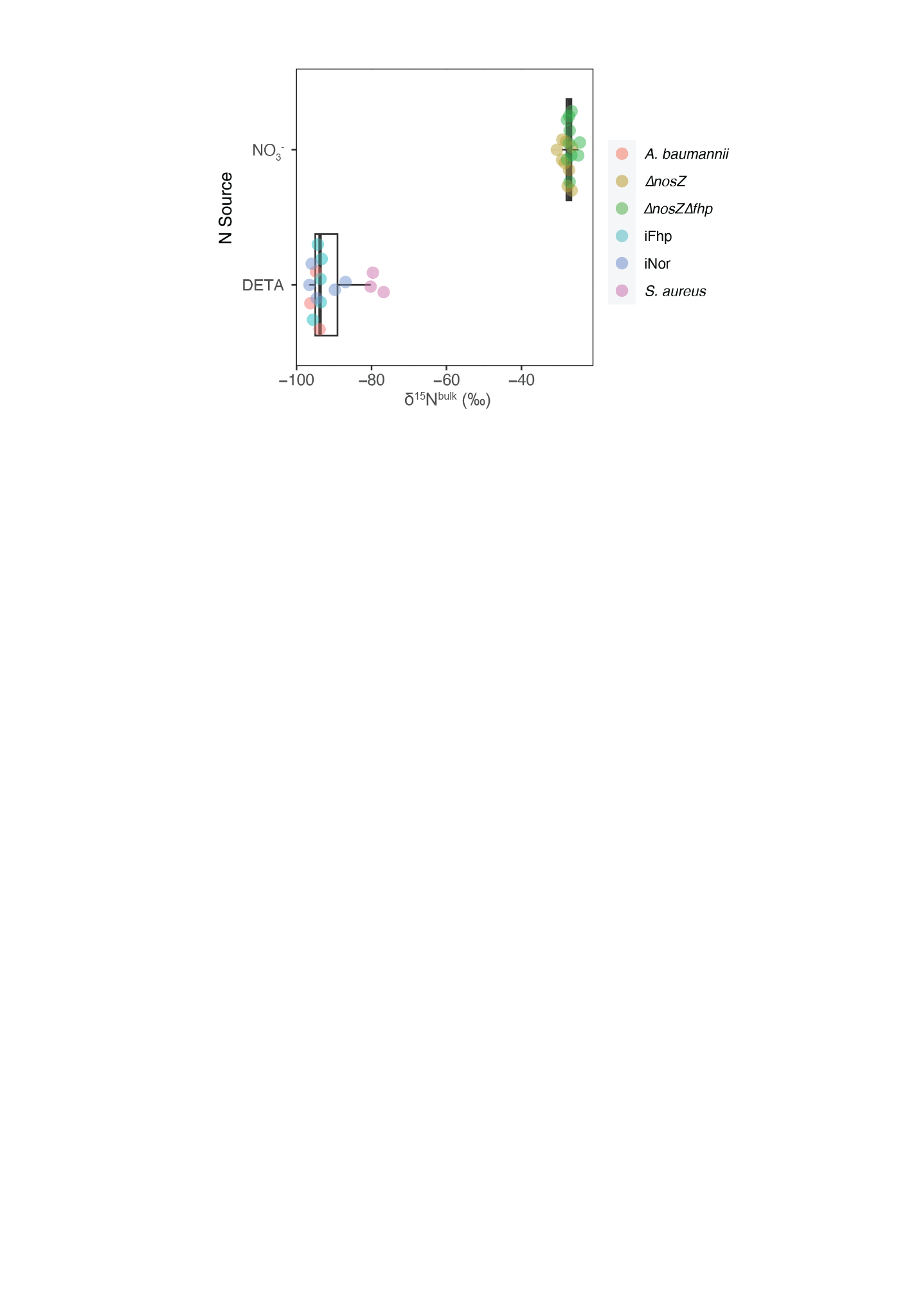

##### Fig S8: δ^15^N^bulk^ for all non-WT *Pa* UCBPP-PA14 experiments.

Data is shown separated by primary NO source – endogenous NO through reduction of nitrate through the denitrification pathway (“NO_3_^-^”), or exogenous NO through addition of DETA NONOate (“DETA”). *A. baumannii*, iNOR, iFhp and *S. aureus* strains (red, navy, cyan, purple; bottom portion of plot, -91.0 ± 6.5‰, mean ± s.d.) were all grown as suspension assays. Though strains were incubated anoxically with both DETA NONOate and nitrate, NO was only sourced from DETA NONOate because nitrate could not be reduced to NO – i.e. native denitrification pathway was deleted in iNOR and iFhp, and does not exist in wild-type *A. baumannii* and *S. aureus* strains. 𝛥*nosZ* and 𝛥*nosZ*𝛥*fhp* strains (green, yellow; top portion of plot, -27.4 ± 1.4‰) were grown as batch cultures and incubated anoxically with only nitrate. See Main Text for strain description and growth conditions.

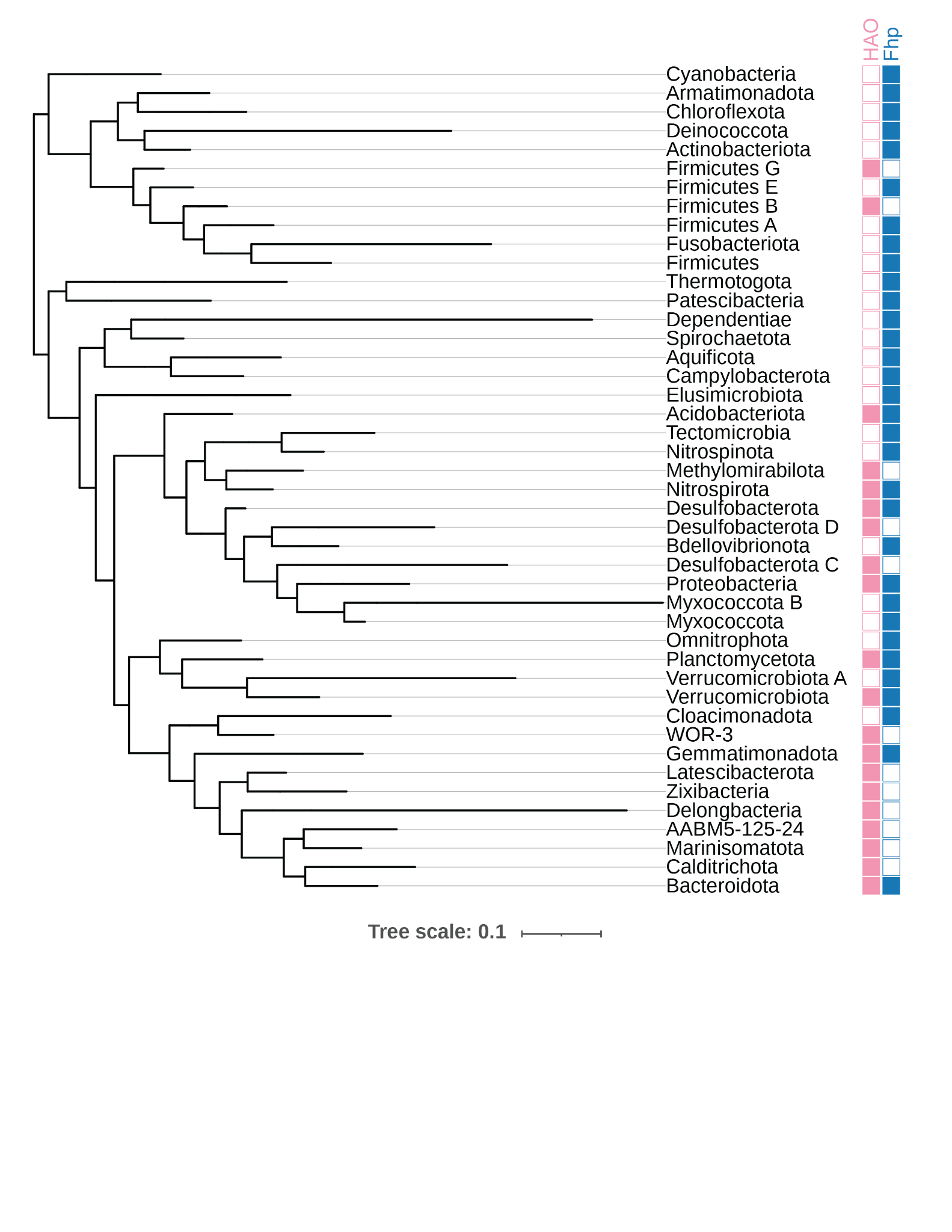

##### Fig. S9: Phylogeny of HAO and Fhp in Bacteria.

Genome hits for hydroxylamine oxidoreductase (HAO; K10535) and Fhp (K05916) in bacteria at the phylum level from AnnoTree [(*27*)](https://sciwheel.com/work/citation?ids=6807209&pre=&suf=&sa=0). Search parameters were used: % identity: 30; E value: 0.00001; % subject alignment: 70; % query alignment: 70. Results were visualized using the Interactive Tree of Life (iTOL); leaves without hits for HAO or Fhp were trimmed for clarity. HAO catalyzes the oxidation of NH_2_OH to NO [(*29*)](https://sciwheel.com/work/citation?ids=7439951&pre=&suf=&sa=0) and is used as a proxy for ammonia oxidizing bacteria.

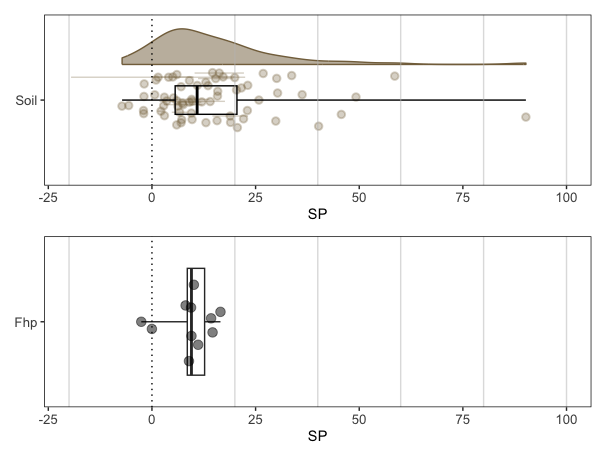

##### Fig. S10: Soil vs. Fhp SP data.

Upper panel shows a density plot of all SP data for soil, with discrete data and boxplot shown below. See Fig. S11 below for references used in literature compilation; each data point represents an individual measurement; for studies that did not report the full dataset, the mean ± s.d. is used instead (shown as a circle with error bar). Boxplot shows five number summary [min, first quartile, median, third quartile, max]. Bottom panel shows boxplot with five number summary for all measurements of Fhp across WT *Pa*, *S. aureus*, and *A. baumannii*. Fhp statistics are [-2.6, 8.5, 9.5, 12.7, 16.5]. Soil statistics are [-7.2, 5.6, 10.9, 20.5, 90.2]. 11.9% of all soil SP data lie within the first and third quartiles (8.5 to 12.7‰) of our measured Fhp data, while 64% lie within the full range (-2.6 to 16.5‰).

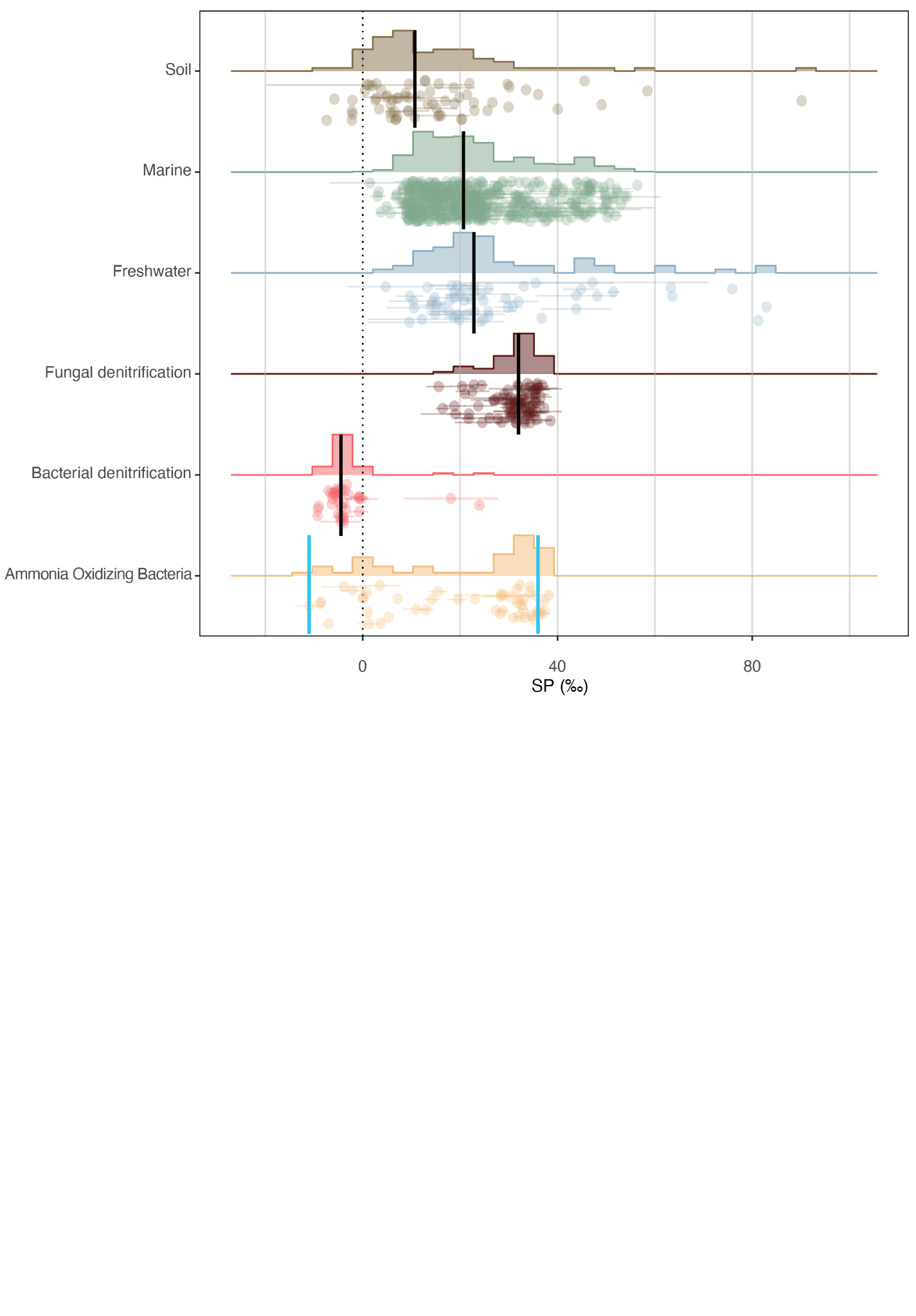

##### Fig. S11. Full literature compilation of environmental and end-member SP values.

Literature compilation of environmental (31 references; *n* = 622[(*21*, *30*–*59*)(*21*, *30*–*59*)](https://sciwheel.com/work/citation?ids=14743626,15212131,15212484,5538575,15212800,1395191,15212851,1254022,15491616,682904,8346772,8346813,8346765,8346759,4708288,15218597,15218614,15218644,15218709,8346796,1145690,1143021,15219125,2244033,15219171,9878514,15219236,2199505,2040354,15219277,15219286&pre=&pre=&pre=&pre=&pre=&pre=&pre=&pre=&pre=&pre=&pre=&pre=&pre=&pre=&pre=&pre=&pre=&pre=&pre=&pre=&pre=&pre=&pre=&pre=&pre=&pre=&pre=&pre=&pre=&pre=&pre=&suf=&suf=&suf=&suf=&suf=&suf=&suf=&suf=&suf=&suf=&suf=&suf=&suf=&suf=&suf=&suf=&suf=&suf=&suf=&suf=&suf=&suf=&suf=&suf=&suf=&suf=&suf=&suf=&suf=&suf=&suf=&sa=0,0,0,0,0,0,0,0,0,0,0,0,0,0,0,0,0,0,0,0,0,0,0,0,0,0,0,0,0,0,0&dbf=0&dbf=0&dbf=0&dbf=0&dbf=0&dbf=0&dbf=0&dbf=0&dbf=0&dbf=0&dbf=0&dbf=0&dbf=0&dbf=0&dbf=0&dbf=0&dbf=0&dbf=0&dbf=0&dbf=0&dbf=0&dbf=0&dbf=0&dbf=0&dbf=0&dbf=0&dbf=0&dbf=0&dbf=0&dbf=0&dbf=0)) and pure culture (15 references; *n* = 172[(*8*, *20*, *22*, *24*, *25*, *60*–*69*)(*8*, *20*, *22*, *24*, *25*, *60*–*69*)](https://sciwheel.com/work/citation?ids=2285408,1158730,1236289,10820078,13028172,15096099,5719748,1635108,1645745,1635109,5384884,1437434,5714832,1480049,2243094&pre=&pre=&pre=&pre=&pre=&pre=&pre=&pre=&pre=&pre=&pre=&pre=&pre=&pre=&pre=&suf=&suf=&suf=&suf=&suf=&suf=&suf=&suf=&suf=&suf=&suf=&suf=&suf=&suf=&suf=&sa=0,0,0,0,0,0,0,0,0,0,0,0,0,0,0&dbf=0&dbf=0&dbf=0&dbf=0&dbf=0&dbf=0&dbf=0&dbf=0&dbf=0&dbf=0&dbf=0&dbf=0&dbf=0&dbf=0&dbf=0) SP data. Environmental data (Soil, Marine, Freshwater) are *in situ* measurements; therefore soil incubation studies were not included. Soil includes forest, cropland, grassland and wetlands. Freshwater includes lakes, rivers and groundwater. To the best of our ability, each data point in the Environmental data represents an individual measurement; for studies that did not report the full dataset, the mean ± s.d. is used instead (shown as circle with error bar). Biogenic endmembers (Fungal denitrification, Bacterial denitrification and Ammonia Oxidizing Bacteria (AOB)) are from *in vitro* studies of pure strains or enzymes. SP values of ammonia oxidizing archaea (AOA) are not included because measurements were performed on enrichment cultures rather than purified strains[(*68*, *70*)(*68*, *70*)](https://sciwheel.com/work/citation?ids=454660,1480049&pre=&pre=&suf=&suf=&sa=0,0&dbf=0&dbf=0), though they have SP values that lie within the positive spread of AOB studies (≈10-30‰). Each data point represents a unique biological replicate; for studies that did not report a full data set, the mean ± s.d. is used instead. Vertical black bar shows median; blue vertical bars of AOB indicate end-member values (roughly -11‰ for nitrifier-denitrification and 36‰ for NH2OH decomposition) that the SP of AOB has been found to vary between based on growth conditions[(*20*)(*20*)](https://sciwheel.com/work/citation?ids=5714832&pre=&suf=&sa=0&dbf=0) due to multiple pathways of N_2_O formation[(*71*)(*71*)](https://sciwheel.com/work/citation?ids=8346774&pre=&suf=&sa=0&dbf=0). All analyses and data visualization were performed using R Statistical Software (v4.1.0;[(*13*)(*13*)](https://sciwheel.com/work/citation?ids=12719082&pre=&suf=&sa=0&dbf=0)) and the ggplot2 package (v3.3.6;[(*28*)(*28*)](https://sciwheel.com/work/citation?ids=14398114&pre=&suf=&sa=0&dbf=0)).

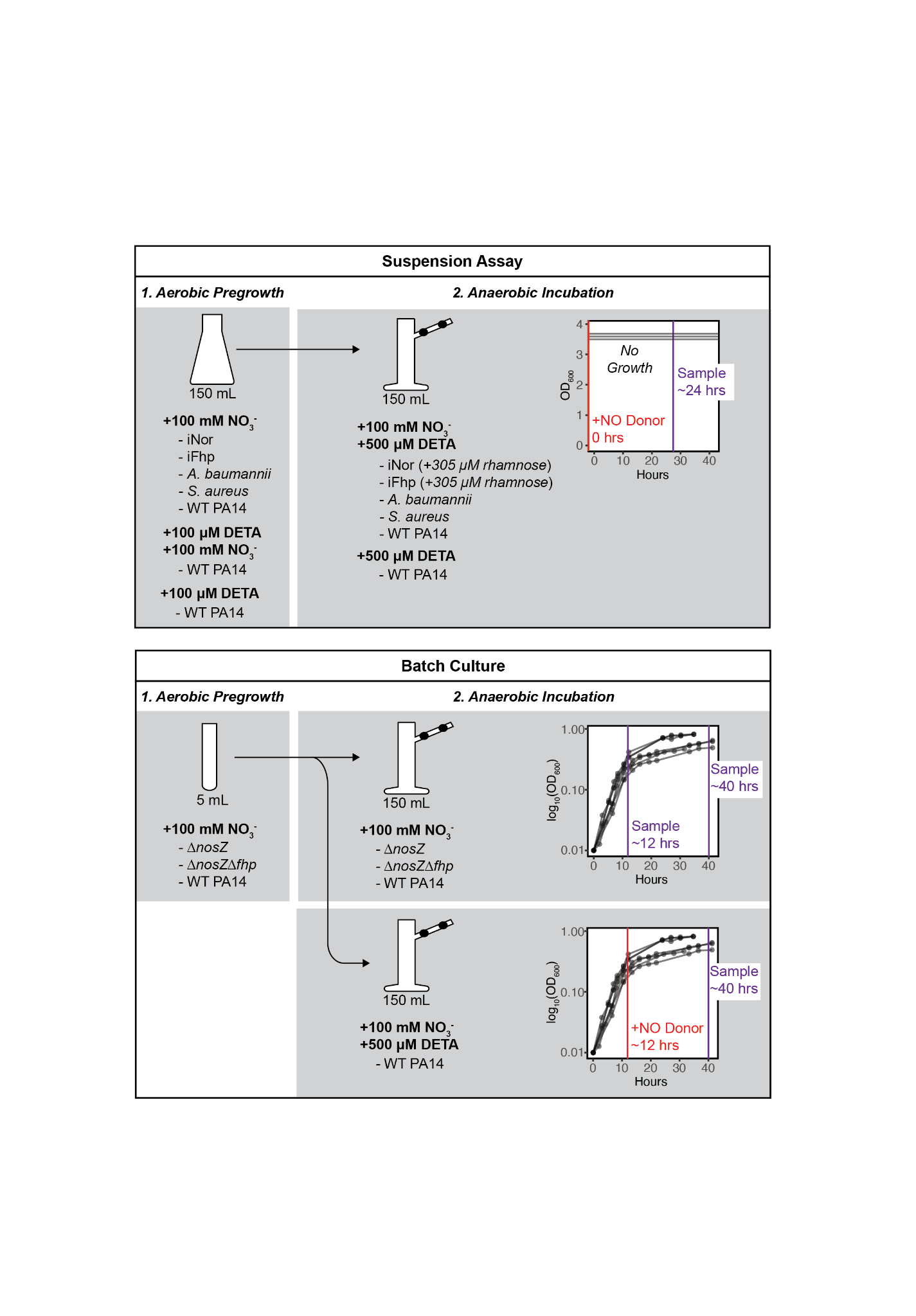

##### Fig. S12. Culturing conditions.

Suspension assays or batch culture assays were performed for this study. Nitrate and / or DETA NONOate was added at varying concentrations for each experiment – concentrations are written in bolded text while strains grown in those conditions are listed below. All strains were first grown in an aerobic pre-growth before anaerobic incubation in the vacuum sampling flask.

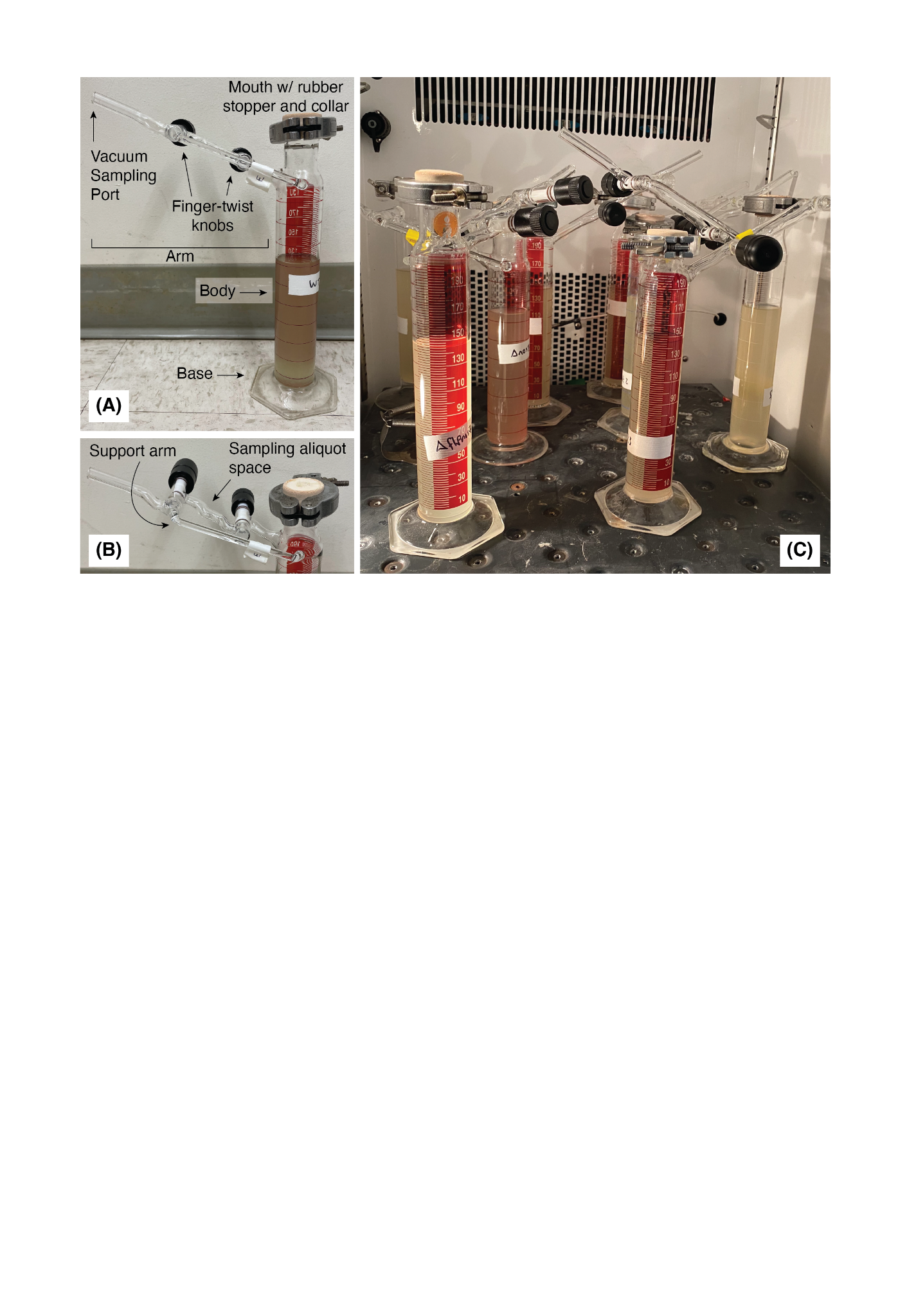

##### Fig. S13. Vacuum sampling flasks for anoxic incubation.

Custom glass vacuum flasks, building off a similar design used in [(*8*, *9*)](https://sciwheel.com/work/citation?ids=5719748,1830225&pre=&pre=&suf=&suf=&sa=0,0), were made in collaboration with the Caltech Glass Shop. **(A, B)** The mouth of a 200 mL borosilicate graduated cylinder was removed and a narrowed neck for a ~2.5 cm diameter rubber stopper was attached. After media and cells were added, a finger-tightened metal collar was placed around the rubber stopper as an additional safeguard. ⅜” gauge glass tubes with two finger-twist knobs were added at the neck of the flask for headspace sampling on the vacuum line. A small sampling aliquot space was retained between the two knobs to isolate gas from the culturing media and the vacuum line. **(C)** Multiple flasks incubated at 37°C, as in a typical sampling workflow. Flasks were incubated in the dark; the light was turned on for the photo.

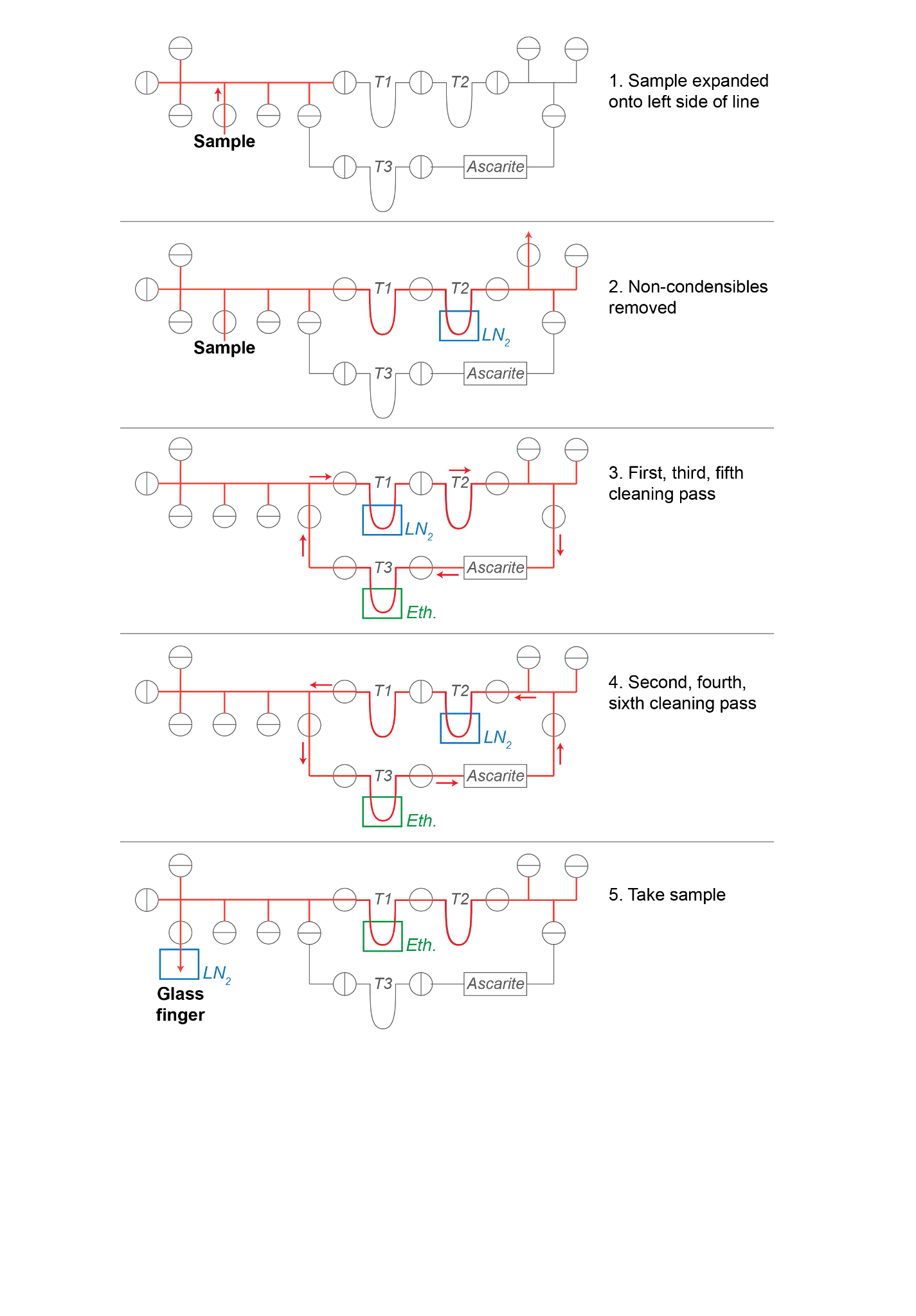

##### Fig. S14. N_2_O Distillation.

Diagram of ultra torr vacuum line used to distill N_2_O from headspace samples. Red indicates portions of the line with sample gas. T1, T2 and T3 refer to different traps that were submerged in either liquid nitrogen (LN_2_) or an ethanol / dry ice slurry (Eth.). Ascarite tube used for CO_2_ removal is shown as a rectangle; valves are shown as circles with the center line indicating if the valve was closed or not; directionality of sample gas flow is shown with red lines.

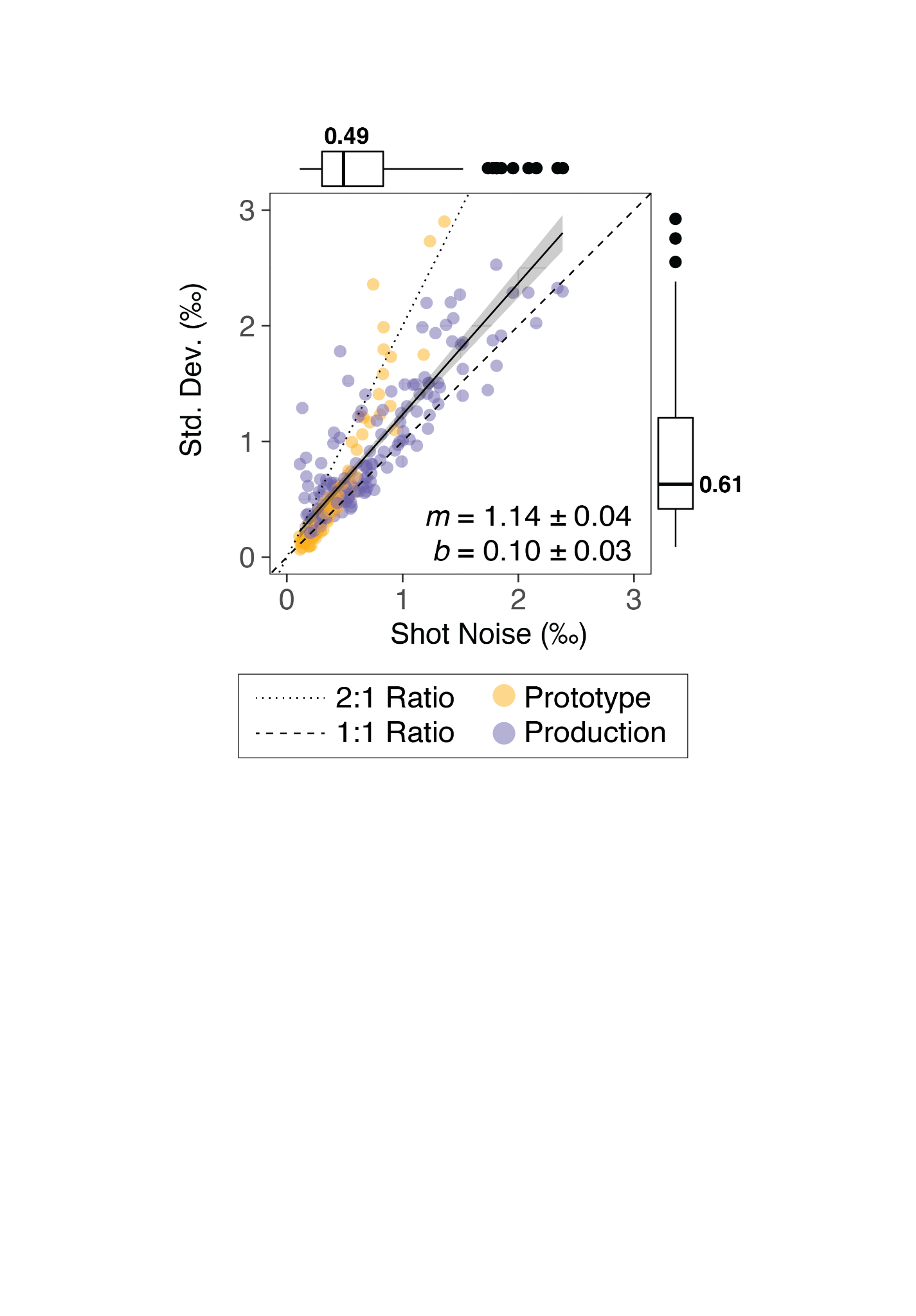

##### Fig. S15. Shot noise and limits of precision.

Calculated shot noise (x-axis) vs. observed standard deviation (y-axis) for all measurements (*n*=79) across the Prototype (yellow) and Production (purple) Ultra for all isotopic measurements. Dashed and dotted lines show 1:1 and 2:1 ratios of standard deviation (std. dev.) to shot noise respectively. Boxplots show median (bolded line and values), upper and lower quartiles (hinges) and 150% of the interquartile range (whiskers); outliers are shown as discrete dots. All analyses and data visualization were performed using R Statistical Software (v4.1.0; R Core Team 2021, [(*13*)](https://sciwheel.com/work/citation?ids=12719082&pre=&suf=&sa=0)) and the ggplot2 package (v3.3.6; Wickham, 2016, [(*28*)](https://sciwheel.com/work/citation?ids=14398114&pre=&suf=&sa=0)).

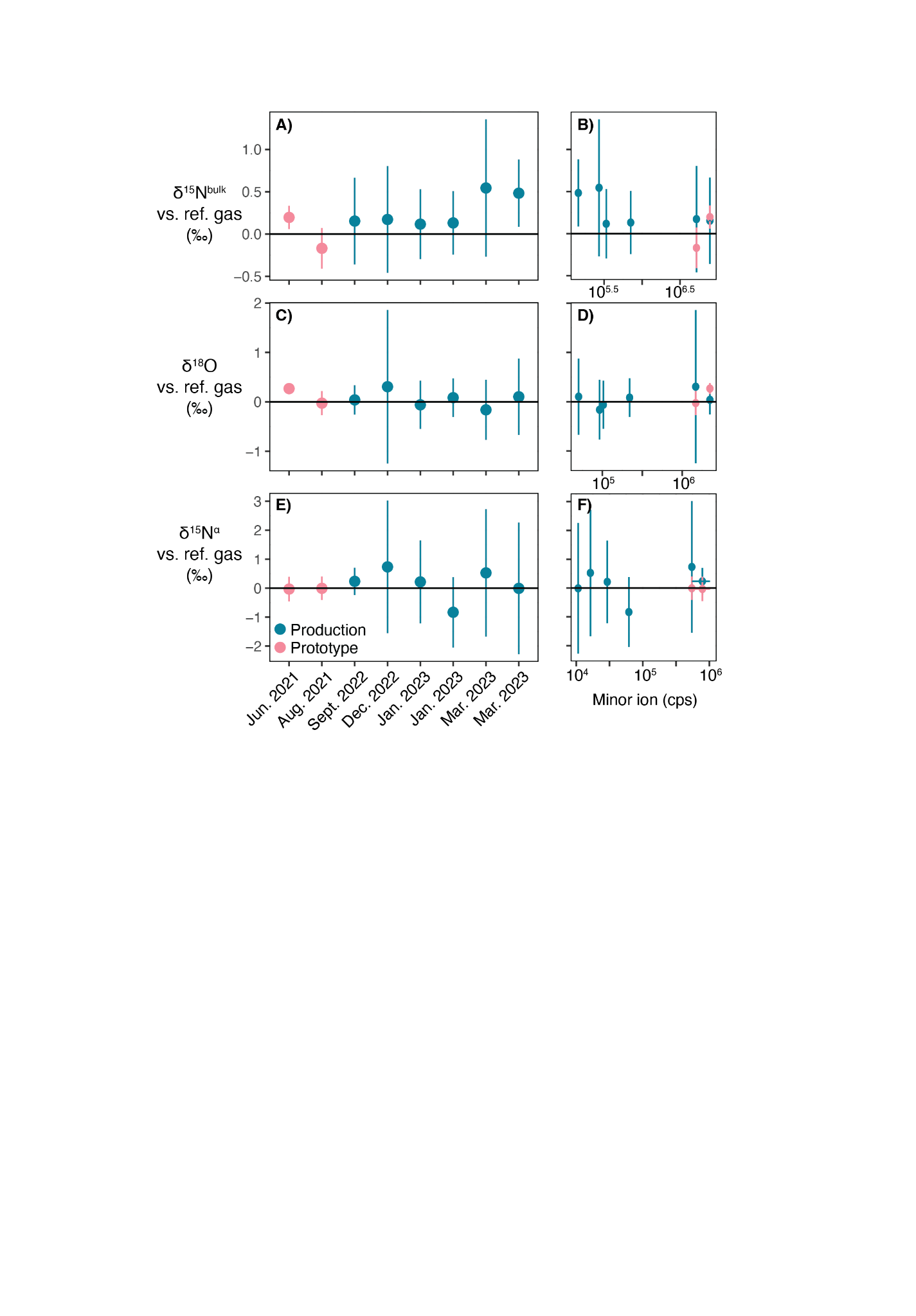

##### Fig. S16. Zero enrichment tests

Zero enrichment test results on the Production (blue) and Prototype Ultras (pink); zero enrichment tests are where the reference gas was measured as a sample against itself. **(A)**, **(C)** and **(E)** show δ^15^N^bulk^, δ^18^O and δ^15^N^ɑ^ vs. experimental session. **(B)**, **(D)** and **(F)** show the same δ values vs. minor ion intensity (cps) for Mass 45, 46 and 31 respectively. All analyses and data visualization were performed using R Statistical Software (v4.1.0; R Core Team 2021, [(*13*)](https://sciwheel.com/work/citation?ids=12719082&pre=&suf=&sa=0)) and the ggplot2 package (v3.3.6; Wickham, 2016, [(*28*)](https://sciwheel.com/work/citation?ids=14398114&pre=&suf=&sa=0)).

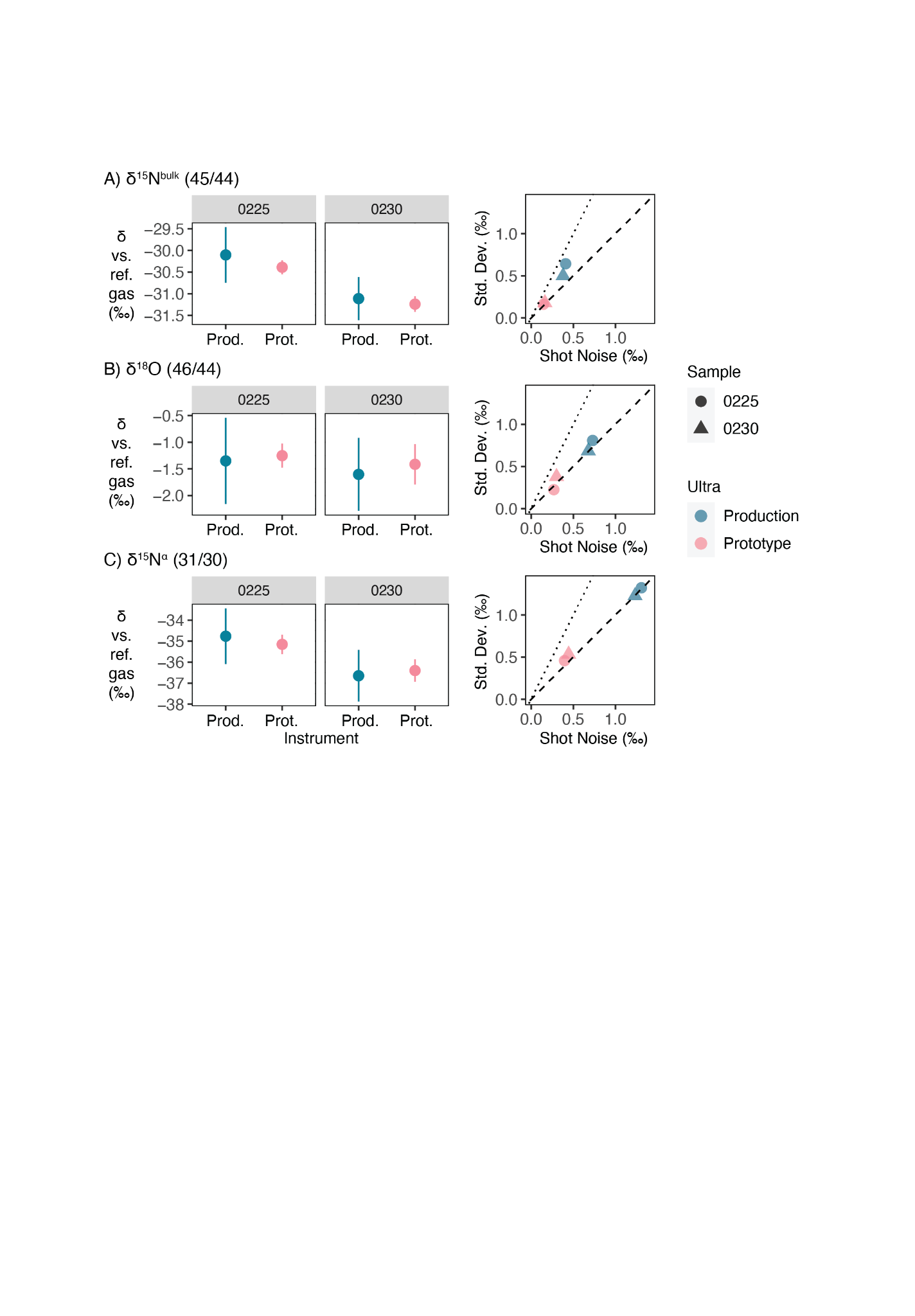

##### Fig. S17. Measurement consistency across instruments.

Two samples, 0225 (circles) and 0230 (triangles), were measured on both the Prototype (pink) and Production (blue) Ultras to gauge measurement accuracy across instruments. Measurements are reported vs. the reference gas for **(A)** δ^15^N^bulk^, **(B)** δ^18^O and **(C)** δ^15^N^ɑ^. Right column shows shot noise (‰) on the x-axis and std. dev. (‰) on the y-axis; 1:1 ratio is shown as a dashed line and 2:1 ratio is shown as a dotted line.

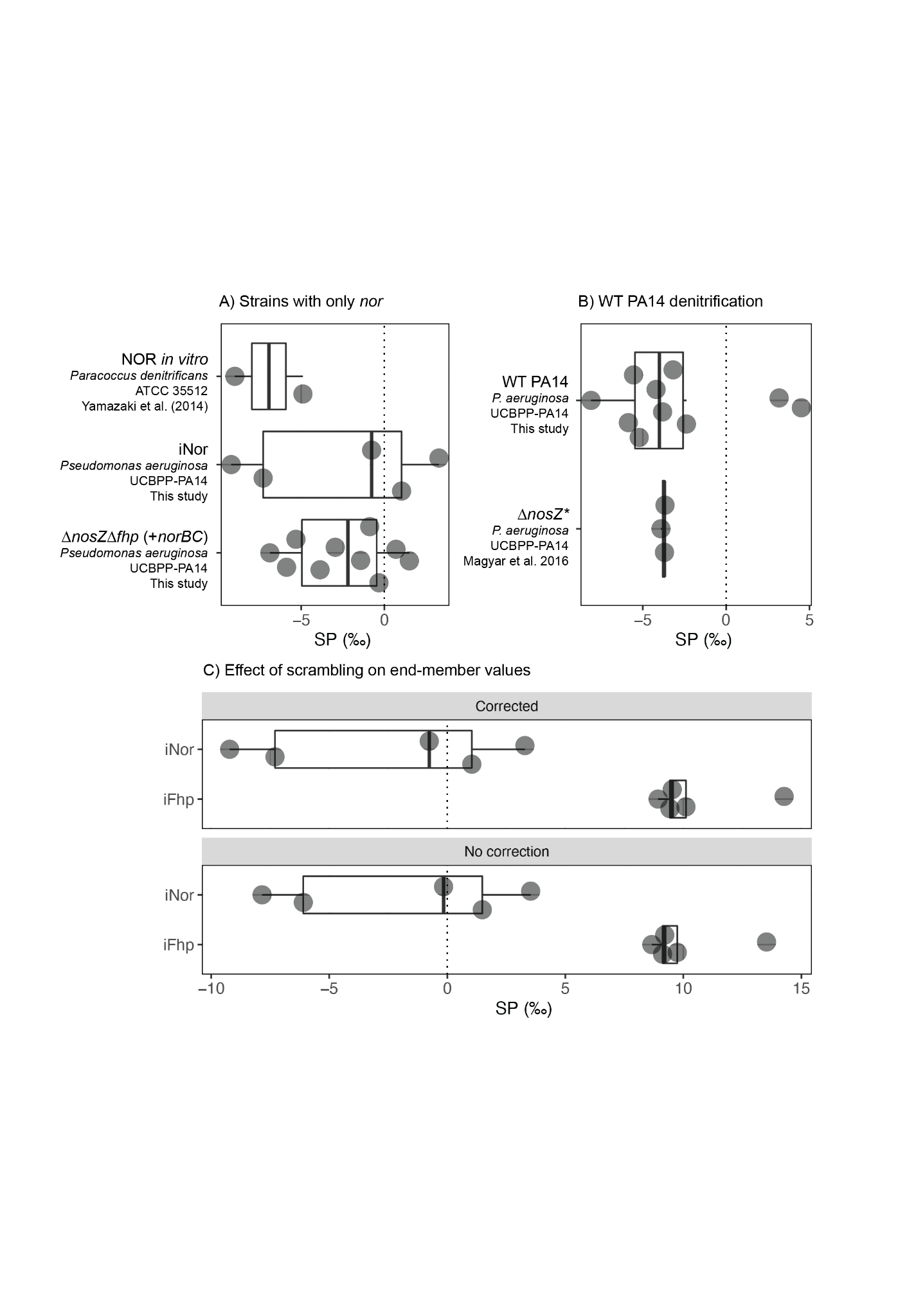

##### Fig. S18. Comparison of scrambling corrected values vs. prior studies.

**(A)** Scrambling-corrected SP values for strains with only NOR (iNor, Δ*nosZ*Δ*fhp*) compared to previously published values by[(*22*)(*22*)](https://sciwheel.com/work/citation?ids=2285408&pre=&suf=&sa=0&dbf=0) of an *in vitro* NOR purified from *Paracoccus denitrificans* ATCC 35512. To accurately compare data across studies, each data point shows one biological replicate; data from[(*22*)(*22*)](https://sciwheel.com/work/citation?ids=2285408&pre=&suf=&sa=0&dbf=0) is presented as the average of Experiments A and the single data point from Experiment C from their study. *ΔnosZΔfhp*, measured on the Prototype Ultra, was corrected using 𝛾 = 0.110 and iNOR, measured on the Production Ultra, was corrected using 𝛾 = 0.045. **(B)** Comparison of scrambling-corrected WT PA14 SP values from this study and[(*9*)(*9*)](https://sciwheel.com/work/citation?ids=1830225&pre=&suf=&sa=0&dbf=0), both grown in similar batch culture, denitrifying conditions. The strain used in[(*9*)(*9*)](https://sciwheel.com/work/citation?ids=1830225&pre=&suf=&sa=0&dbf=0) was reported as “*Pseudomonas aeruginosa* strain PA14 Δ*nosZ*,” but we found through PCR amplification of the *nosZ* gene location that the deletion was not successful – therefore, this strain is the same as our study and is the WT strain. This is indicated as “Δ*nosZ**” in the figure. Values from this study and[(*9*)(*9*)](https://sciwheel.com/work/citation?ids=1830225&pre=&suf=&sa=0&dbf=0) were both measured on the Prototype Ultra and corrected using 𝛾 = 0.110.

SUPPLEMENTARY TABLES

| **Species** | **SP References** | **Fhp accession number** | **NOR accession number** |
| --- | --- | --- | --- |
| *Pseudomonas aeruginosa* | This study; Magyar et al. (2016); Magyar et al. (2017) | A0A0H2ZC95 | A0A0H2ZLE2 (NorB); A0A0H2ZKE8 (NorC) |
| *Pseudomonas fluorescens* | Toyoda et al. (2005) | A0A448BJZ8 or A0A8H2RPK4 | A0A0D0T5F4 (NorB); A0A0D0S4Z1 (NorC) |
| *Paracoccus denitrificans* | Toyoda et al. (2005); Ostrom et al. 2007 | A1B2P2 | Q51663 (NorB); Q51662 (NorC) |
| *Pseudomonas chlororaphis*; *Pseudomonas aureofaciens* subsp. nov., comb. nov. | Magyar et al 2017; Kantnerova et al 2022; Sutka et al. 2006; Haslun et al. 2018 | A0A5M7CAB6 | Q9F0W6 (NorB); Q9F0W7 (NorC) |
| *Pseudomonas* *stutzer*i (*Stutzerimonas stutzeri*) | Ostrom et al 2007 | Q5W5T4 | P98008 (NorB); Q52527 (NorC) |

##### Table S1. Fhp and NOR accession numbers for previously measured bacterial denitrifiers.

The accession number for Fhp or NorB and NorC of denitrifying strains used in prior SP studies. A close strain relative, whose genome has been sequenced, was used. Fhp is also annotated as Hmp or NOD (nitric oxide dioxygenase). [(*69*)](https://sciwheel.com/work/citation?ids=2243094&pre=&suf=&sa=0) used "*Pseudomonas stutzeri* (provided by J. M. Tiedje)" and *Pseudomonas denitrificans* ATCC 13867; *P. stutzeri* is also known as *Stutzerimonas stutzeri*. [(*8*, *9*)](https://sciwheel.com/work/citation?ids=1830225,5719748&pre=&pre=&suf=&suf=&sa=0,0) used *Pseudomonas aeruginosa* UCBPP-PA14 and *Pseudomonas aureofaciens* ATCC 13985. [(*60*)](https://sciwheel.com/work/citation?ids=1158730&pre=&suf=&sa=0) used *Pseudomonas fluorescens* ATCC 13525 and *Paracoccus denitrificans* ATCC 17741 (also known as 19376). [(*24*)](https://sciwheel.com/work/citation?ids=13028172&pre=&suf=&sa=0) used *Pseudomonas aureofaciens* ATCC 13985. [(*25*)](https://sciwheel.com/work/citation?ids=1437434&pre=&suf=&sa=0) used *Pseudomonas aureofaciens* ATCC 13985 and *Pseudomonas chlororaphis* ATCC 43928. However, DNA-DNA hybridization experiments has led to the reclassification of *P. aureofaciens* into *P. chlororaphis* [(*72*)](https://sciwheel.com/work/citation?ids=13028211&pre=&suf=&sa=0) – therefore the strain “*Pseudomonas aureofaciens* ATCC 13985” used by [(*8*, *24*, *25*)](https://sciwheel.com/work/citation?ids=1437434,13028172,5719748&pre=&pre=&pre=&suf=&suf=&suf=&sa=0,0,0) is now a subspecies of *P. chlororaphis* with the proposed taxonomic name “*P. chlororaphis* subsp. *aureofaciens* subsp. nov., comb. nov. [with the type strain DSM 6698^T^ (=ATCC 13985^T^=NCIMB 9030^T^)] [(*72*)](https://sciwheel.com/work/citation?ids=13028211&pre=&suf=&sa=0).” Therefore, *P. aureofaciens* and *P. chlororaphis* are grouped together in the graph above. [(*63*)](https://sciwheel.com/work/citation?ids=15096099&pre=&suf=&sa=0) specified that they use the strains “*Pseudomonas chlororaphis* subsp. chlororaphis (ATCC 43928; *P. chlororaphis*) and *Pseudomonas chlororaphis* subsp. *aureofaciens* (ATCC 13985; *P. aureofaciens*).”

| **Bacterial Phylum** | **Number of genome hits** | **Proportion of all hits** | **Number of genomes in clade** |
| --- | --- | --- | --- |
| Proteobacteria | 3761 | 52.90% | 9474 |
| Myxococcota | 28 | 0.39% | 168 |
| Myxococcota_B | 1 | 0.01% | 2 |
| Bdellovibrionota | 4 | 0.06% | 110 |
| Desulfobacterota | 4 | 0.06% | 560 |
| Nitrospirota | 3 | 0.04% | 138 |
| Nitrospinota | 1 | 0.01% | 22 |
| Tectomicrobia | 1 | 0.01% | 4 |
| Acidobacteriota | 14 | 0.20% | 380 |
| Bacteroidota | 318 | 4.47% | 3781 |
| Gemmatimonadota | 4 | 0.06% | 101 |
| Cloacimonadota | 1 | 0.01% | 27 |
| Verrucomicrobiota | 36 | 0.51% | 478 |
| Verrucomicrobiota_A | 2 | 0.03% | 52 |
| Planctomycetota | 52 | 0.73% | 376 |
| Omnitrophota | 2 | 0.03% | 83 |
| Elusimicrobiota | 3 | 0.04% | 66 |
| Campylobacterota | 60 | 0.84% | 323 |
| Aquificota | 4 | 0.06% | 39 |
| Spirochaetota | 17 | 0.24% | 310 |
| Dependentiae | 1 | 0.01% | 26 |
| Patescibacteria | 13 | 0.18% | 1131 |
| Thermotogota | 1 | 0.01% | 63 |
| Firmicutes | 1086 | 15.28% | 2737 |
| Fusobacteriota | 3 | 0.04% | 70 |
| Firmicutes_A | 62 | 0.87% | 2636 |
| Firmicutes_E | 2 | 0.03% | 39 |
| Actinobacteriota | 1492 | 20.99% | 4261 |
| Deinococcota | 10 | 0.14% | 92 |
| Chloroflexota | 31 | 0.44% | 520 |
| Armatimonadota | 1 | 0.01% | 36 |
| Cyanobacteria | 91 | 1.28% | 727 |
| Total: | 7109 | 1 | 28832 |

##### Table S2. Fhp AnnoTree query results in Bacteria.

Fhp (KEGG ID K05916) query results in AnnoTree [(*27*)](https://sciwheel.com/work/citation?ids=6807209&pre=&suf=&sa=0) at the phylum level for Bacteria. Default search parameters were used: % identity: 30; E value: 0.00001; % subject alignment: 70; % query alignment: 70.

| **Bacterial Phylum** | **Number of genome hits** | **Proportion of all hits** | **Number of genomes in clade** |
| --- | --- | --- | --- |
| Proteobacteria | 7 | 0.4375 | 9474 |
| Verrucomicrobiota | 3 | 0.1875 | 478 |
| Planctomycetota | 6 | 0.375 | 376 |
| Total: | 16 | 1 | 10328 |

##### Table S3: Fhp and HAO AnnoTree query results in Bacteria.

Fhp (K05916) and HAO (K10535) query results in AnnoTree [(*27*)](https://sciwheel.com/work/citation?ids=6807209&pre=&suf=&sa=0) at the phylum level for Bacteria. HAO was used as a proxy for ammonia oxidizing bacteria (AOB). Default search parameters were used: % identity: 30; E value: 0.00001; % subject alignment: 70; % query alignment: 70.

| **Species** | **Putative Yhb/Fhp Accession** | **% Yhb (identical/similar)** | **% Fhp (identical/similar)** |
| --- | --- | --- | --- |
| *Aspergillus tenneri* | XP_033421839 | 36.8/55.4 | 40.8/59.9 |
| *Fusarium oxysporum f. sp. rapae* | KAG7407474 | 38.0/53.1 | 41.1/58.2 |
| *Fusarium zealandicum* | KAF4977403 | 38.1/52.2 | 39.9/56.8 |
| *Metarhizium anisopliae* | KAF5132848 | 35.3/51.0 | 36.1/53.4 |
| *Penicillium cf. griseofulvum* | KAJ5211151 | 36.4/54.8 | 41.3/59.2 |
| *Trichoderma arundinaceum* | RFU80218 | 38.4/52.8 | 36.4/52.3 |

##### Table S4. Predicted occurrence of Yhb/Fhp orthologues in p450nor-containing fungi.

Representative fungal species with p450nor-attributed N_2_O SP values [(*64*, *65*)](https://sciwheel.com/work/citation?ids=1635108,1645745&pre=&pre=&suf=&suf=&sa=0,0) were obtained from NCBI, and pairwise sequence analysis was performed against *S. cerevisiae* Yhb (Accession NP_011750) and *P. aeruginosa* UCBPP-PA14 Fhp with EMBOSS Needle [(*26*)](https://sciwheel.com/work/citation?ids=12816401&pre=&suf=&sa=0).

| Primer ID | Sequence | Description |
| --- | --- | --- |
| fhp_1-55 | TCTGCAGGAATTCCTCGAGAAGCTTATGTTGTCCAATGCCCAACGTGCC | Amplify *fhp* for generation of iFhp strain, forward |
| fhp_1-56 | GCAAGGCCTTCGCGAGGTACCTCAGGCGTCCAGCGCGGC | Amplify *fhp* for generation of iFhp strain, reverse |
| norCBD_2-5 | TCTGCAGGAATTCCTCGAGAAGCTTATGTCCGAGACCTTTACCAAAGGCATGGC | Amplify *nor* for generation of iNOR strain, forward |
| norCBD_2-6 | GCAAGGCCTTCGCGAGGTACCTCAGCGGCGCAGGCGCCG | Amplify *nor* for generation of iNOR strain, reverse |
| fhp DN F | GCATGCGTCAGGAGTCATCTTGGACGCCTGAAGCGACGGG | Amply flanking region of *fhp* for mutagenesis |
| fhp DN R | CATGATTACGAATTCGAGCTAGCACGCAGCCCAGCAGGAT | Amply flanking region of *fhp* for mutagenesis |
| fhp UP F | ACGACGGCCAGTGCCAAGCTTGGCCGAACAATTCGCTTTC | Amply flanking region of *fhp* for mutagenesis |
| fhp UP R | CCCGTCGCTTCAGGCGTCCAAGATGACTCCTGACGCATGC | Amply flanking region of *fhp* for mutagenesis |
| fhp Genotyping F | GCAAGGGATTGGTGGTCATTTCG | Sequencing/confirming *fhp* deletion |
| fhp Genotyping R | CATCAGCCTGGAACGATCAAGC | Sequencing/confirming *fhp* deletion |

##### Table S5. Primers used in this study.

Primers used for amplification and deletion of *fhp* and *norCBD* in parent strain *Pseudomonas aeruginosa* UCBPP-PA14.

| **Sample ID** | **Sample Description** | **Moles of N_2_O sampled** | **OD_600_** | **Corrected N_2_O / OD_600_** |
| --- | --- | --- | --- | --- |
| BLK 1 | DETA NONOate only | 4.7E-08 | *NA* | *NA* |
| BLK 2 | DETA NONOate only | 5.6E-08 | *NA* | *NA* |
| Sa 1 | *Staphylococcus aureus* | *NA* | 2.979 | *NA* |
| Sa 2 | *Staphylococcus aureus* | 7.4E-07 | 3.986 | 5.80E+06 |
| Ab 1 | *Acinetobacter baumannii* | 1.4E-06 | 2.794 | 2.00E+06 |
| Ab 2 | *Acinetobacter baumannii* | 1.2E-06 | 3.546 | 2.99E+06 |

##### Table S6. Results of N_2_O screen for Fhp-only strains.

Sample pressures from the direct-injection bellows of the Thermo Scientific Ultra High-Resolution Isotope Ratio Mass Spectrometer (HR-IRMS); moles of N_2_O was calculated using the ideal gas law. OD_600_ indicates optical density at 600 nm. Corrected N_2_O indicates that moles of N_2_O have been corrected for the moles of N_2_O produced in the no-cell controls (BLK 1 and 2). A concentration of 500 𝜇M was used for DETA NONOate.

| **Batch** | **Material** | **δ^15^N (‰)** | **Moles of N added** |
| --- | --- | --- | --- |
| Feb102022 | Nitrate | 0.15 ± 0.26 | 0.015 |
| Feb102022 | SCFM Amended | -0.82 ± 0.19 | 0.00420 |
| Aug302021 | Nitrate | 1.79 ± 0.02 | 0.015 |
| Aug302021 | SCFM Amended | -1.77 ± 0.13 | 0.00420 |
| Aug192021 | Nitrate | -0.73 ± 0.08 | 0.0350 |
| Aug192021 | SCFM Amended | -1.91 ± 0.14 | 0.00420 |

##### Table S7. Batch culture nitrogen substrate.

Batch culture experiments were carried out using three batches of nitrate and SCFM Amended media – Feb102022, Aug302021 and Aug192021. δ^15^N values (mean ± s.d.) are corrected for tin capsule blanks; moles of N indicate total how many moles of N from nitrate or SCFM Amended were added to the total 150 mL culture volume. Additional moles of nitrate were accidentally added in the Aug192021 batch.

|  | **0100** | **0101** | **0112** |
| --- | --- | --- | --- |
| δ^15^N^bulk^ | 0.12 ± 0.32‰ | 0.07 ± 0.21‰ | 0.41 ± 0.42‰ |
| δ^18^O | -0.03 ± 0.30‰ | 0.02 ± 0.41‰ | -2.25 ± 0.90‰ |
| δ^15^N^⍺^ | -0.09 ± 0.28‰ | -0.39 ± 0.47‰ | -0.34 ± 1.17‰ |

##### Table S8. Distillation and vacuum flask blanks

Isotopic measurements of N_2_O distillation blank (0100, 0101) and no-cells vacuum flask blank (0112). Values (mean ± s.d.) are reported relative to the working reference gas, Caltech Ref Gas.

| Measurement | Value vs. intl. reference |
| --- | --- |
| δ^15^N^bulk^ | 4.21‰ |
| δ^15^N^⍺^ | 7.53‰ |
| δ^15^N^𝜷^ | 0.89‰ |
| δ^15^N^SP^ | 6.64‰ |
| δ^18^O | 39.96‰ |

##### Table S9. Caltech Ref Gas

Caltech Ref Gas, the working reference gas used in this study, reported relative to the international standards of Air for N and VSMOW for O. Caltech Ref Gas was previously characterized by colleagues at Tokyo Tech [(*8*, *9*)](https://sciwheel.com/work/citation?ids=5719748,1830225&pre=&pre=&suf=&suf=&sa=0,0). Reference gas was obtained through Matheson Gas at Ultra High Purity (99.99%).

| Value | RM5 Reported | RM5_1 Measured | RM5_2 Measured |
| --- | --- | --- | --- |
| δ^15^N^bulk^ | 33.44 ± 0.05‰ | 34.26 ± 0.36‰ | 34.01 ± 0.53‰ |
| δ^15^N^⍺^ | 43.54 ± 0.91‰ | 42.80 ± 1.27‰ | 42.54 ± 1.99‰ |
| δ^15^N^𝜷^ | 23.34 ± 1.29‰ | 25.72 ± 1.32‰ | 25.48 ± 2.06‰ |
| δ^15^N^SP^ | 20.2 ± 0.91‰ | 17.08 ± 1.83‰ | 17.06 ± 2.86‰ |
| δ^18^O | 39.52 ± 0.15‰ | 39.31 ± 0.36‰ | 39.41 ± 0.48‰ |

##### Table S10. Characterization of RM5.

Values for RM5 are taken from[(*17*)(*17*)](https://sciwheel.com/work/citation?ids=14308013&pre=&suf=&sa=0&dbf=0). RM5_2 was measured three months after RM5_1 and at a lower sample amount, which caused RM5_2 to have larger measurement uncertainties overall. All values are reported as mean ± s.d. and versus AIR.

| Value | Vs. Caltech Ref Gas | Vs. Intl (No scrambling corr.) | Final reported values |
| --- | --- | --- | --- |
| δ^15^N^bulk^ | -97.39 ± 0.18‰ | -93.59 ± 0.18‰ | -93.59 ± 0.18‰ |
| δ^15^N^⍺^ | -95.53 ± 0.51‰ | -88.72 ± 0.51‰ | -88.54 ± 0.51‰ |
| δ^15^N^𝜷^ | -99.25 ± 0.54‰ | -98.46 ± 0.54‰ | -98.65 ± 0.54‰ |
| δ^15^N^SP^ | 1.86 ± 0.74‰ | 4.87 ± 0.74‰ | 10.11 ± 0.54‰ |
| δ^18^O | -16.58 ± 0.25‰ | 22.72 ± 0.25‰ | 22.72 ± 0.25‰ |

##### Table S11. Example of scrambling correction.

Values are reported as mean ± s.e. for one measurement of iFhp on the Production Ultra using g = 0.045. The raw measurement (“Vs. Caltech Ref Gas”) is first corrected to the international standard of AIR or VSMOW (“Vs. Intl (No scrambling corr.”), and then the scrambling correction is applied (“Final Reported Values,” Eqn. S12).

| **Sample ID (# of N)** | **δ^15^N (‰)** |
| --- | --- |
| Full Donor (5 N) | -22.95 ± 0.15‰ |
| Decomposed Donor (3 N) | -23.54 ± 0.24‰ |
| Released N (2 N) | -22.08 ± 0.29‰ |

##### Table S12. Isotopic composition of DETA NONOate

Measured δ^15^N values of the full and decomposed NO-donor, DETA NONOate. The δ^15^N of the released nitrogens was calculated by mass balance (Eqn. S13). Values represent mean ± s.d. of three replicates.

| **Bacterial Species** | **NCBI Fhp Reference Sequence** |
| --- | --- |
| *Pseudomonas aeruginosa* UCBPP-PA14 | WP_003138913.1 |
| *Pseudomonas cavernae* | AYC30948.1 |
| *Stenotrophomonas maltophilia* | KAF1052024.1 |
| *Pseudomonas toyotomiensis* | WP_206418516.1 |
| *Acinetobacter baumannii* | SST07660.1 |
| *Klebsiella pneumoniae* | SVJ66665.1 |
| *Methyloversatilis discipulorum* | WP_019918880.1 |
| *Azotobacter chroococcum* | WP_131300631.1 |
| *Escherichia coli* | MRF39747.1 |
| *Paraburkholderia sp. SOS3* | SOS3WP_075159372.1 |
| *Bacillus sp. TH86* | TH86MBK5304199.1 |
| *Streptococcus pneumoniae* | CJK98816.1 |
| *Bacillus cereus* group | WP_103625387.1 |
| *Yersinia pseudotuberculosis* | WP_038401225.1 |
| *Burkholderia cepacia complex* | WP_080747301.1 |
| *Staphylococcus aureus* | WP_072399307.1 |

##### Table S13. Sequence identifiers used for Fig S1 phylogeny.
